# Sertad1 is elevated and plays a necessary role in synaptic loss, neuron death and cognitive impairment in a model of Alzheimer’s disease

**DOI:** 10.1101/2024.08.06.606751

**Authors:** Naqiya Ambareen, Kusumika Gharami, Subhas C. Biswas

**Author notes:** Correspondence should be addressed to Subhas C. Biswas, Cell Biology and Physiology Division, CSIR-Indian Institute of Chemical Biology, 4 Raja S. C. Mullick Road, Kolkata 700 032, India.

## Abstract

Dysfunctional autophagy is a primary characteristic of Alzheimer’s disease (AD) pathogenesis. How autophagic impairment leads to cellular changes that contributes to AD pathogenesis remains unclear. To study this further, we assessed levels of autophagy related proteins in 5xFAD mice brain at different ages and found their robust upregulation in cortex and hippocampus suggesting increased induction of autophagy with disease progression but failed clearance. We have identified a transcriptional coregulator Sertad1, as a key mediator of dysfunctional autophagy in AD mice. We found a progressive elevation in Sertad1 levels in 5xFAD mice with age compared to wild-type (WT) mice. Sertad1 knockdown in 5xFAD mice brain lowered levels of autophagy related proteins and lysosome marker, LAMP1 suggesting its role in autophagy flux modulation. FoxO3a is an important transcriptional regulator of the autophagy network and lies at the nexus of autophagy-apoptosis cross-talk. We found that Sertad1 knockdown blocked nuclear translocation of FoxO3a along with a restoration in Akt activity. Further, we showed that knockdown of Sertad1 in 5xFAD mice brain improved cognitive functions along with a remarkable restoration in synaptic health and dendritic spine density. Taken together, our results demonstrated that autophagy is robustly induced with disease progression but it is impaired; Sertad1 knockdown restored autophagy defects, synaptic loss and improved learning and memory in AD models. Thus, we propose that Sertad1 acts in a multimodal manner regulating crucial cell death pathways including apoptosis and autophagy and could be an excellent target for therapeutic intervention to combat a multifactorial disorder such as AD.

## Introduction

Alzheimer’s disease (AD) is the leading cause of dementia accounting for 60-80% of the total number of dementia cases with no cure available yet [1]. It is a multi-factorial progressive disorder that imposes severe burden on caregivers. Hardy and Higgins first proposed the amyloid cascade hypothesis in 1992, which places amyloid-β (Aβ) accumulation as the central tenet of AD pathogenesis and other events such as aberrant tau phosphorylation and accumulation as neurofibrillary tangles, neuroinflammation, synaptic dysfunction and neuron loss occur downstream of Aβ accumulation [2]. Many studies support this neuron-centric linear hypothesis of AD pathology [3–7]. Contrarily, many groups have raised a debate over the linearity of the amyloid cascade as it fails to address the quiet incubation time of AD [8]. The ‘biochemical phase’ of AD is marked by the accumulation of amyloid plaques and neurofibrillary tangles that exerts immense proteopathic or aggregate stress disrupting the normal signalling functions of the brain [8, 9]. The biochemical phase paves way for the ‘cellular phase’ of the disease characterized by chronic inflammation, circuitry imbalances and cell death. Failure in the cellular mechanisms to maintain homeostasis culminates in the ‘clinical phase’ resulting in clinical manifestation of dementia [8]. The autophagy-lysosomal network (ALN), specifically macroautophagy, referred to as autophagy from now on, is a primary modulator of the proteopathic stage of AD pathogenesis [10]. Mysterious double membrane vesicles that were later identified as immature autophagic vacuoles (AVs) containing undigested protein aggregates and other cellular waste were reported to accumulate within swollen dystrophic neurites in the AD brain [10–12]. Although many studies have pointed out that there are defects in ALN, the temporal profiling of autophagy with disease progression has not been done yet. Other studies have shown that disruptions in the synapse and decline in hippocampal LTP precede Aβ plaque deposition [13, 14]. Plaque deposition starts at around 3-4 months in 5xFAD mice [15, 16] that further contributes to impairment in several cellular processes exacerbating pathogenic changes in a feed-forward mechanism.

Studies suggest that apoptosis and autophagy have some common inducers and an intricate cross-talk exists between these two processes [17]. Previously, our group has reported that both autophagy and apoptosis are simultaneously induced in AD models [18, 19]. Sertad1, also known as Trip-Br1 and Sei1 plays an important role in apoptotic cell death in a Nerve Growth Factor (NGF) deprivation and DNA damage model of neurodegeneration [20]. Sertad1 was previously known to be involved in cell-cycle processes, as a regulator of CDK4 activity through its binding and subsequent activation [21]. Additionally, Sertad1 has been implicated in various cancers [22–24]. As a transcriptional co-regulator, it is known to stimulate transcription factor activities of E2F1 [25], p53 [26] and SMAD1 [27] in diverse models. Sertad1 may be regarded as a master regulator as it binds to several proteins including CDK4 [21], XIAP [28], E2F1/DP1 [25] and PP2A [29], thus regulating several cellular signalling processes. It was reported to be elevated in an ischemic stroke model and activated the CDK4/p-Rb pathway, thereby contributing to neuronal injury [30]. We found few reports that provide evidence that Sertad1 might be involved in the autophagy-lysosomal pathway in breast cancer cells [22, 24] but there is no evidence of the involvement of Sertad1 in autophagy modulation in neurodegenerative diseases. Current research shows that a single unwavering approach directly targeting Aβ plaques is not sufficient to fight a multifactorial disorder such as AD.

In this study, we showed that autophagy-related and lysosomal proteins are upregulated in 5xFAD mice with disease progression indicating increased induction of autophagy in response to aggregate stress. We are the first to identify the involvement of a transcriptional coactivator, Sertad1 in autophagy modulation in AD pathogenesis. We reported an age-dependent increase of Sertad1 in the AD transgenic mice brain and its correlation with autophagic dysfunction. We discovered that Sertad1 regulates autophagy through the Akt/FoxO3a axis. We discovered that Sertad1 knockdown mice showed improved cognitive behaviour in a series of behavioural paradigms that we assessed along with amelioration of synapse loss observed in AD pathology. Taken together, our study sheds light on the mechanism of autophagy modulation by Sertad1 and highlights its role as an excellent target for therapeutic intervention.

## Results

### Autophagy is robustly induced in 5xFAD mice brain but fails to reach completion

Autophagy in neurons is tightly regulated in a spatio-temporal manner and is responsible for maintaining homeostasis [31, 32]. However, in neurodegenerative disorders such as AD, autophagy flux is impaired, resulting in accumulation of indigested autophagosomes within dystrophic neurites present in the AD patients’ brain [11, 33, 34]. Here, we have performed a detailed characterization of autophagy and lysosome proteins at different ages of AD transgenic mice (5xFAD) as compared to its corresponding age-matched WT mice. Details of animal model used in this study is given in the Materials and Methods section. Since cortex and hippocampus are most affected areas in AD associated memory impairment [35], we have isolated cortex and hippocampus tissues from 1 month, 3 months, 6 months and 12 months old 5xFAD mice along with their corresponding age-matched WT mice. LC3 and p62 are common markers of autophagy flux. Cytosolic LC3-I is lipidated by its conjugation with phosphatidylethanolamine (PE) to form LC3-II that is present on autophagosome membrane. The expression level of LC3-II and LC3-I to LC3-II conversion as indicated by their ratio, gives a clear indication of the number of autophagosomes [36, 37]. Here, we found that LC3-II is significantly upregulated in 5xFAD mice in cortical and hippocampal lysates of 3 months, 6 months and 12 months of age as compared to their respective age-matched WT mice (Figure 1A, 1B, 1C, 1H, 1I and 1J). p62/SQSTM1 is an autophagy receptor protein which links autophagy with the ubiquitin-proteosomal system. p62 can recognize and interact with K63 linked ubiquitinated proteins and possesses LC3-interacting domain (LIR) via which it targets selective cargoes to the autophagosomes [38]. Defects in autophagy flux result in p62 accumulation due to impaired clearance [37]. We have found significantly increased accumulation of p62 in both cortex and hippocampus of 5xFAD mice at 6 months and 12 months of age (Figure 1A, 1D, 1H and 1K) as compared to their age-matched WT mice. ATG12 conjugates with ATG5 with the help of E1 and E2-like enzymes: ATG7 and ATG10. This complex in turn binds with ATG16L1 to form ATG12-ATG5-ATG16L1 complex which associates with the autophagosome membrane. This complex further helps in LC3 recruitment to the autophagosome and aids in autophagosome elongation [39]. We assessed levels of elongation markers, ATG5 and ATG7 and found a significant increase in their levels at 3 months, 6 months and 12 months of age in the cortex (Figure 1A, 1E and 1F) and hippocampus (Figure 1H, 1L and 1M) in 5xFAD mice as compared to their age-matched WT mice. We further checked the levels of lysosomal marker protein, LAMP1, to assess the accumulation of autolysosomes. We found a significant increase in LAMP1 protein levels in the cortex (Figure 1A and 1G) and hippocampus (Figure 1H and 1N) of 5xFAD mice progressively as compared with their corresponding age-matched WT mice. This data indicates that autophagosomes and lysosomes are robustly synthesized in response to aggregate stress in the AD brain but the system is overburdened resulting in neurotoxic accumulation of undigested autophagosomes and autolysosomes. However, the exact mechanism of autophagy dysfunction, its primary mediators and its role in AD pathogenesis mains elusive.

**Figure 1:**
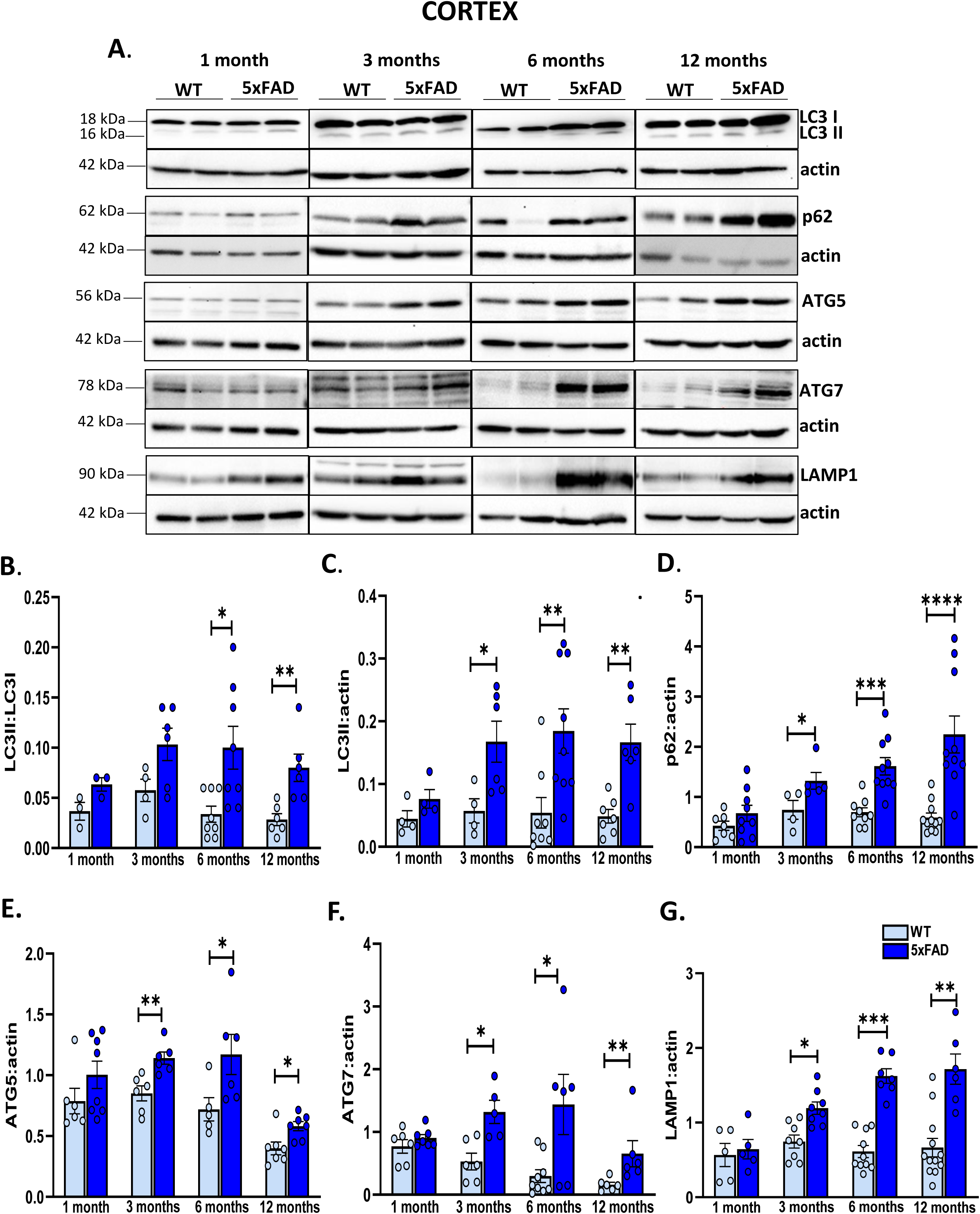
Autophagosomes markers are upregulated in the 5xFAD mice brain. Cortical and hippocampal tissues were isolated from the brains of 5xFAD mice of 1 month, 3 months, 6 months and 12 months old along with their age-matched WT mice, protein lysates were prepared and subjected to Western blotting. Immunoblots showing expression of LC3, p62, ATG5, ATG7 and LAMP1 in the cortex (**A**) and hippocampus (**H**), respectively, of 5xFAD mice at different ages compared with their age-matched WT mice as indicated. Actin was used as loading control. Graphical representation of the relative expression of LC3II:LC3I (**B**), LC3II (**C**), p62 (**D**), ATG5 (**E**), ATG7 (**F**) and LAMP1 (**G**) respectively normalized with actin in cortex. Graphical representation of the relative expression of LC3II:LC3I (**I**), LC3II (**J**), p62 (**K**), ATG5 (**L**), ATG7 (**M**) and LAMP1 (**N**) normalized with actin in hippocampus of 5xFAD mice compared to WT. N= 8-10 mice per group. Data represent Mean±SEM. Statistical analysis was done using two-tailed unpaired T-test, *p<0.05, **p<0.01, ***p<0.001 and ****p<0.0001.

### Sertad1 is elevated in primary cortical neurons upon Aβ treatment and in 5xFAD mice age-dependent manner

We have shown that autophagy and apoptosis are simultaneously induced in response to Aβ treatment indicating an intricate cross-talk between the two cell death modalities [18, 19]. Microarray screening of altered genes in DNA damage induced neurodegeneration revealed a 46-fold increase in the level of Sertad1 and established its role in mediating neuronal cell death in response to DNA damage, NGF deprivation and Aβ exposure [20]. Sertad1 also known as Trip-Br1 and Sei1 is a transcriptional co-regulator that interacts with the PHD bromodomain containing transcription factors [25]. First, we determined the expression levels of Sertad1 in primary cortical neuron cultures upon 1.5 µM Aβ (1-42) treatment at different time points. We observed a significant increase in Sertad1 protein levels at 16 hours and 24 hours of Aβ treatment (Figure 2A and 2B). Next, we isolated and prepared cortical and hippocampal lysates from 1 month, 3 months, 6 months and 12 months old 5xFAD mice along with their age-matched WT mice and examined the expression level of Sertad1 protein. We found a progressive and significant upregulation of Sertad1 in 5xFAD cortex and hippocampus starting at 3 months of age when compared with their age matched WT mice (Figure 2C-2E). We prepared tissue slices from 6 months old 5xFAD mice brain and performed immunofluorescence studies. Immunostaining with Sertad1 and MAP2 antibodies showed that Sertad1 is expressed in neurons and Sertad1 expression is significantly upregulated in the hippocampus of 6 months old 5xFAD mice as compared to their age-matched WT mice (Figure 2F and 2G). Taken together, increase in Sertad1 levels occurs simultaneously with aberrant autophagy induction in 5xFAD mice. We were thus interested to explore the role of Sertad1 in modulating autophagy and AD related cognitive decline.

**Figure 2:**
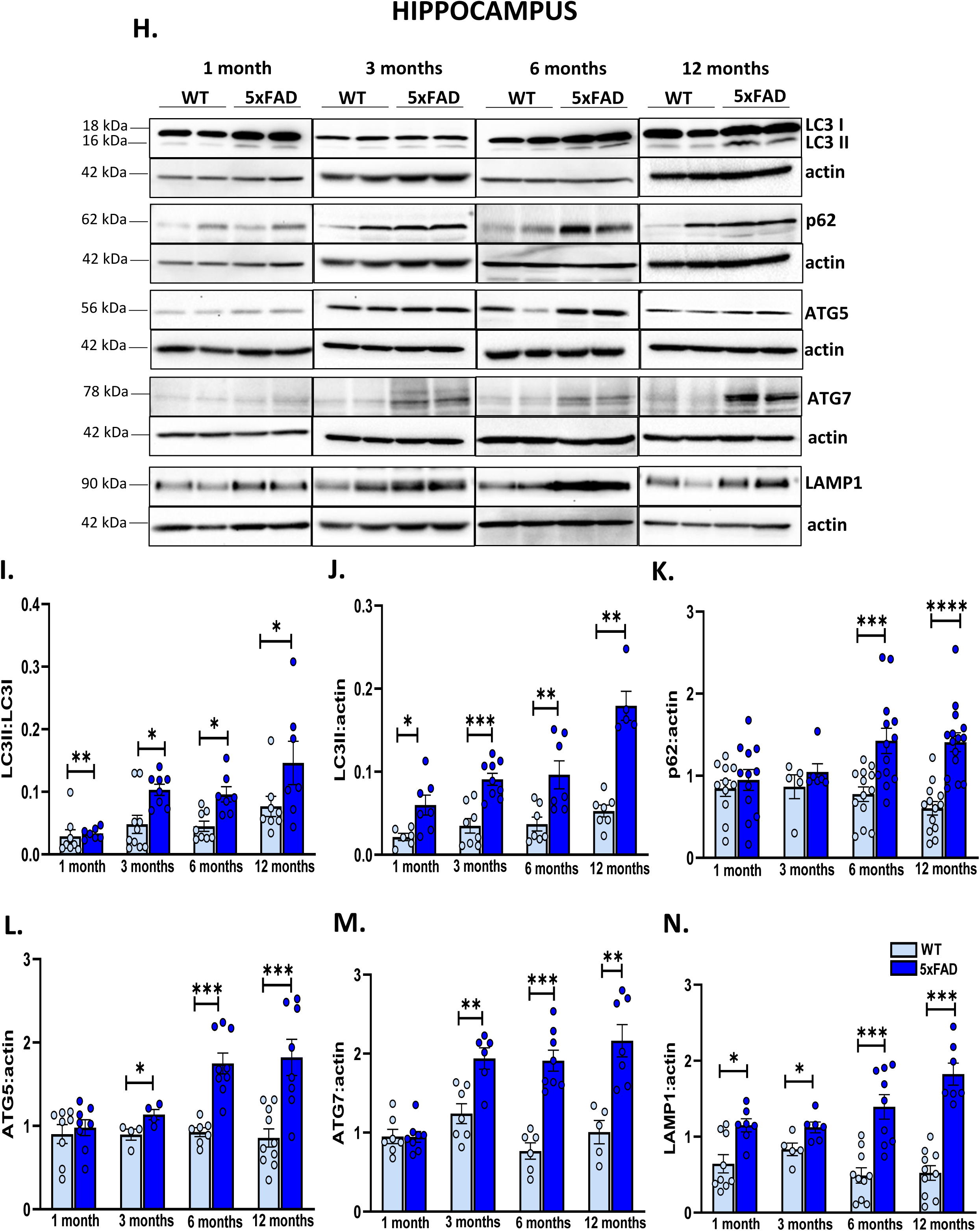
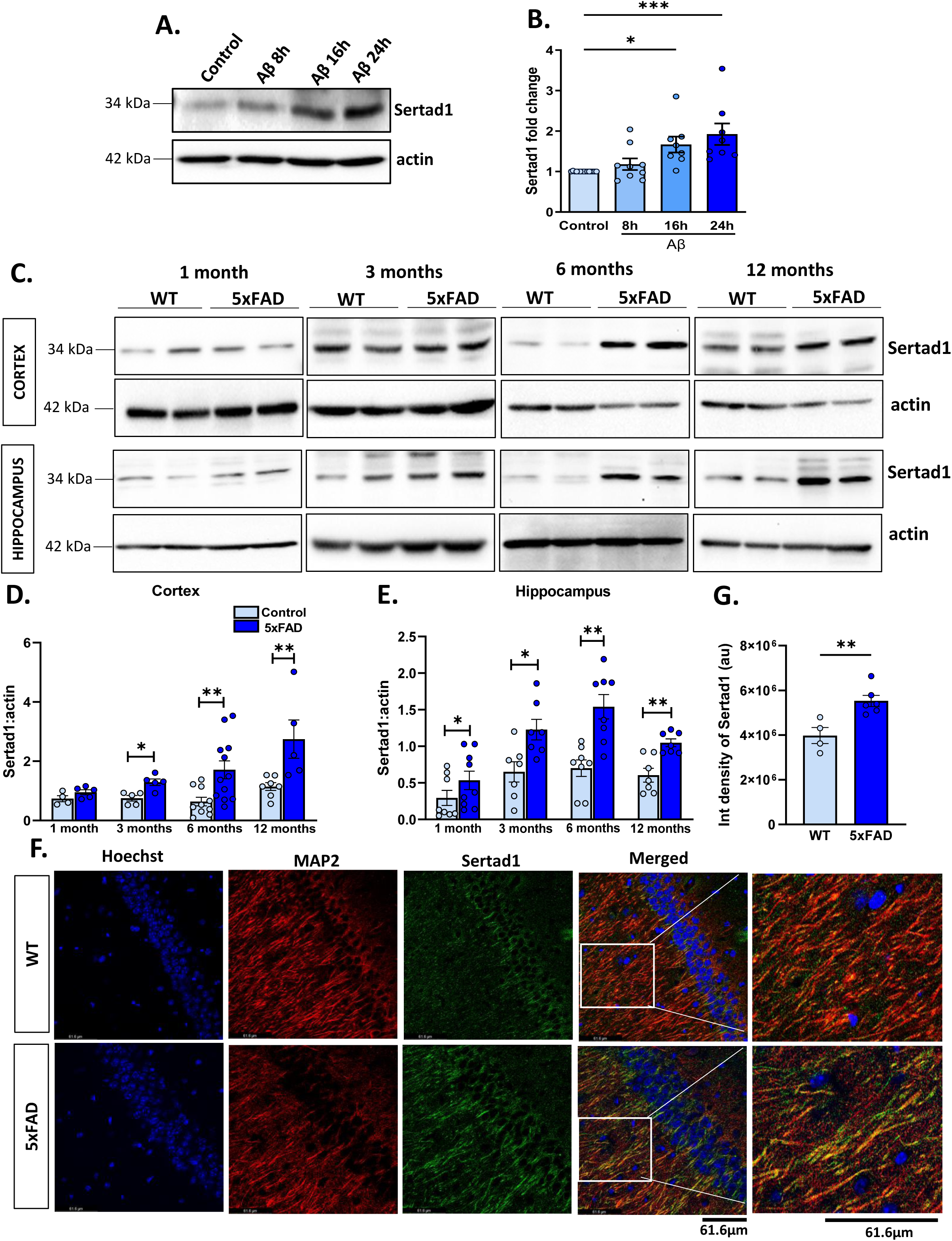
Sertad1 is elevated in primary cortical neurons in response to Aβ and in cortex and hippocampus of 5xFAD mice. Primary cortical neurons were treated with 1.5 µM Aβ for the indicated time points, total cell protein lysates were prepared and subjected to Western blotting. (**A**) Immunoblot showing Sertad1 expression along with the corresponding actin at the indicated time points. (**B**) Graphical representation of fold change of Sertad1 by densitometric analysis using NIH-ImageJ software. (**C**) Whole cell cortical and hippocampal lysates from isolated brains of 1 month, 3 months, 6 months, and 12 months old 5xFAD mice and their corresponding age-matched WT mice were prepared and subjected to Western blotting. Immunoblot showing Sertad1 expression levels in cortex and hippocampus at indicated ages. Actin was used as loading control. Graphical representation of normalized Sertad1:actin in cortex (**D**) and hippocampus (**E**) of 5xFAD mice compared to their age-matched WT mice (N=8 animals/group). (**F**) Expression level of Sertad1 was analysed in coronal brain sections (thickness: 30µm) obtained from 6 months old 5xFAD mice and its age-matched control. Brain sections were co-immunostained with Sertad1 (green) and MAP2 (red) antibodies. Nuclei were stained using Hoechst (blue). Representative image of one of the brain sections is depicted here (magnification: 63X). Difference in Sertad1 expression between 5xFAD and WT mice brain sections was quantified using NIH-ImageJ software. (**G**) Graphical representation of integrated density value of Sertad1 expression (N=3 animals/group). Data represent Mean±SEM. Statistical analysis was done using two-tailed unpaired T-test, asterisks denote statistically significant differences from the corresponding controls; *p<0.05, **p<0.01 and ***p<0.001.

### Knockdown of Sertad1 using shRNA expressing lentiviral particles in 5xFAD mice brain

To decipher the mechanism by which Sertad1 promotes neurodegeneration, we have delivered lentivirus expressing shSertad1 to knockdown Sertad1 in the 5xFAD mice brain. Detailed information about plasmids used, procedure for lentivirus preparation and stereotaxic injection is given in the Materials and Methods section. Firstly, we prepared lentiviral particles expressing shSertad1 and empty vector (EV) using HEK293T cells and validated Sertad1 knockdown by transducing lentivirus in HEK293T cells. 48 hours post-transduction, we observed GFP expression and significantly low expression of Sertad1 expression in shSertad1 transduced HEK293T cells (Figure 3A and 3B). We have chosen 5 months old mice to perform our experiments as cognitive deficiencies become apparent in 5xFAD mice at that age [13, 16]. After validation of Sertad1 knockdown in HEK cells, we injected lentiviral particles expressing shSertad1 and EV in the hippocampus of 5xFAD mice along with age-matched WT mice (coordinates are mentioned in Materials and Methods section). We have shown the experimental design as a schematic diagram in figure 3C. We have found GFP expression both in the cortex and hippocampus of 5xFAD and WT mice infused with shSertad1 and EV (Figure 3D). We validated GFP expression of 5xFAD mice infused with shSertad1 and EV by DAB staining after 14 days of lentiviral particles infusion and observed that lentiviral particles are expressed throughout the brain (Figure 3E). Next, we performed co-immunostaining of Sertad1 and GFP, and found significant downregulation in Sertad1 levels in both hippocampus (Figure 3F and 3G) and cortex (Figure 3H and 3I) of shSertad1 or EV injected 5xFAD mice and WT mice. We have used Sertad1 knockdown 5xFAD mice, EV injected 5xFAD mice along with their corresponding age-matched WT mice for further studies on the role of Sertad1 in disease pathology.

**Figure 3:**
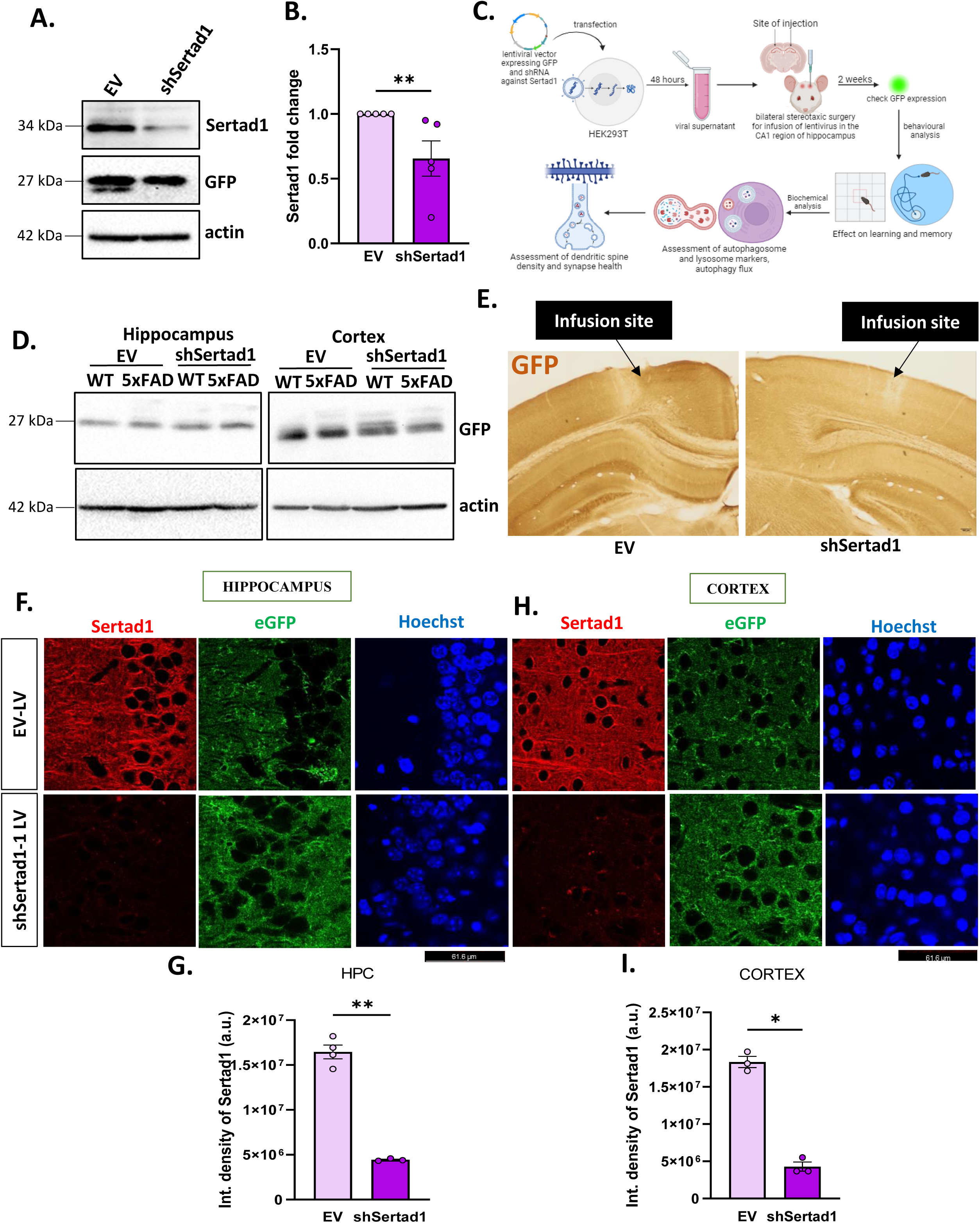
Lentivirus mediated downregulation of Sertad1 in 5xFAD mice. Lentiviral particles expressing shSertad1 alongside EV containing eGFP reporter was prepared as discussed in materials and methods section. HEK293T cells were transduced with shSertad1 and EV expressing lentiviral particles and subjected to Western blotting. (**A**) Representative immunoblot showing Sertad1 and GFP expression is depicted. Actin was used as loading control. (**B**) Graphical representation of fold change of Sertad1 by densitometric analysis using NIH-ImageJ software. (**C**) Schema depicting the plan of work including lentivirus preparation, stereotactic infusion, behavioural analysis and biochemical assays for mechanistic studies. (**D**) Immunoblot showing GFP expression in cortex and hippocampus isolated from mice brain of 14 days post infusion of lentiviral particles. (**E**) Representative image of brain sections stained with anti-GFP antibody using DAB staining protocol depicting the infusion site (magnification: 10X). (**F and H**) Brain sections were co-immunostained with anti-Sertad1 (red) and EGFP (green). Nuclei were stained with Hoechst (blue). Left panel shows the cortex region and right panel shows the CA1 region of the hippocampus. Graphical representation of integrated density value of Sertad1 expression quantified using ImageJ in the hippocampus (**G**) and cortex (**I**) respectively (N=3 animals/group). Data represent Mean±SEM. Statistical analysis was done using two-tailed unpaired T-test, asterisks denote statistically significant differences from the corresponding controls; *p<0.05 and **p<0.01.

### Sertad1 knockdown is protective against Aβ insult, reduces apoptosis levels and amyloid plaque burden in 5xFAD mice

Since Sertad1 elevation coincides with the onset of AD pathology and it is also associated with cell death, we hypothesized that downregulation of Sertad1 using shRNA to be neuroprotective. We have transfected primary cortical neurons with shSertad1-zs-green or shRand-zsgreen (scrambled RNA), maintained them for 48 hours and then treated them with 1.5 µM Aβ. We then tracked the number of live green transfected neurons by monitoring their morphology as well as number at different time points. We observed that shSertad1 transfected neurons retained neuronal morphology and showed higher percentage survival even after 48 hours of Aβ exposure (Figure 4A and 4B). TUNEL assay used to quantify and visualize apoptotic cells [40]. We sacrificed the shSertad1-eGFP and EV-eGFP injected 5xFAD and WT mice after 21 days post-surgery and assessed the percentage of TUNEL positive cells. We found that percentage of TUNEL positive cells was significantly high in 5xFAD mice brain sections compared to WT mice. However, upon Sertad1 knockdown, percentage of TUNEL positive cells significantly reduced in 5xFAD mice brain (Figure 4C and 4D). Amyloid plaque deposition is the primary hallmark of AD pathology and is observed in 5xFAD mice at early stages [15]. We assessed for Aβ plaque burden, number of plaques and amyloid integrated density and found it to be significantly reduced in shSertad1 injected 5xFAD mice as compared to 5xFAD infused with EV [Figure 4E and 4F (i)-(iii)]. These results show that Sertad1 is associated with AD pathogenesis and its knockdown increases neuron survival and reduces number of TUNEL positive cells. Furthermore, lowering Sertad1 levels in 5xFAD mice reduced amyloid pathology in 5xFAD mice. Our next aim was to establish whether there is any link between dysfunctional autophagy and Sertad1.

**Figure 4:**
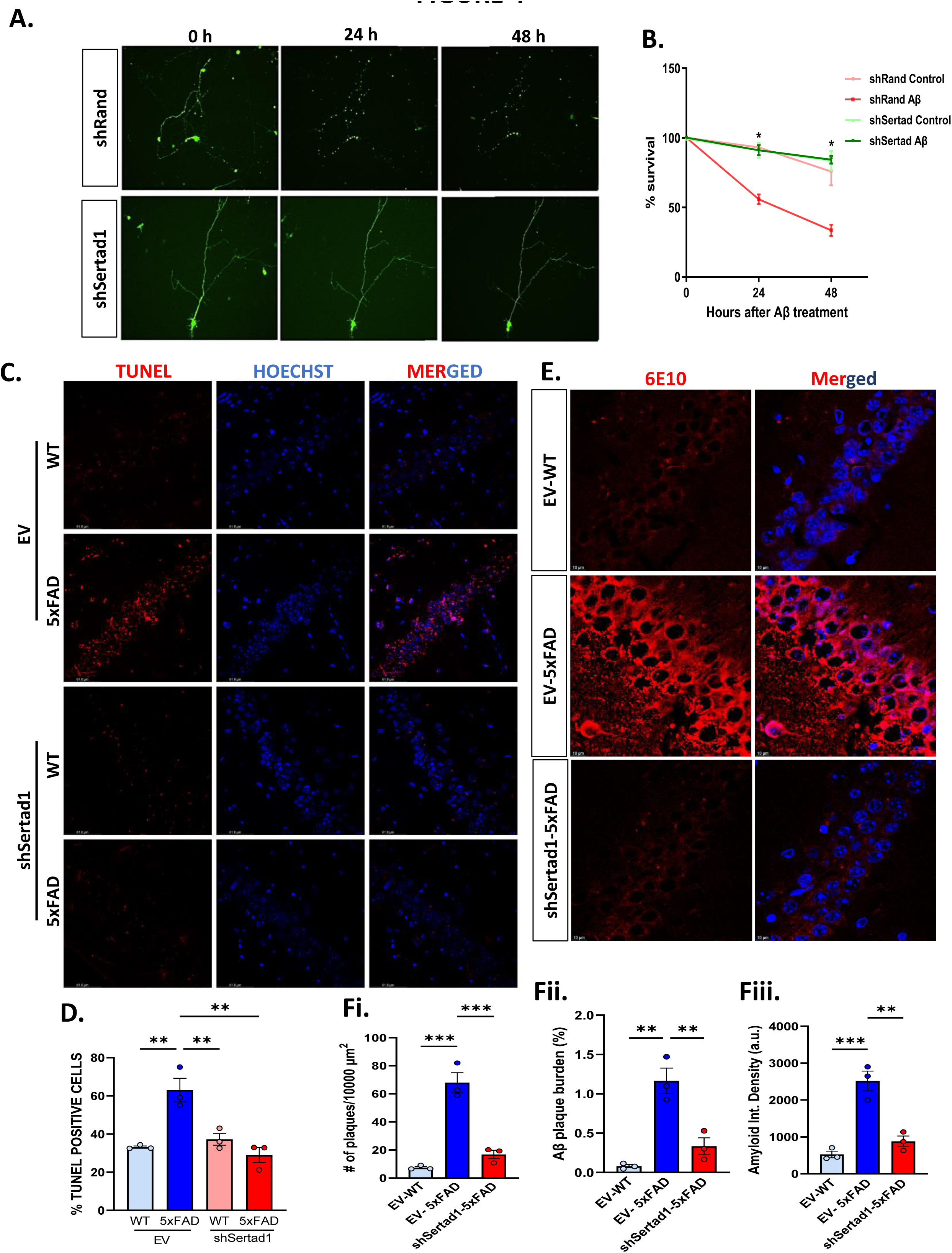
Sertad1 knockdown confers neuroprotection in in-vitro and in-vivo AD models. (**A**) Primary rat cortical neurons were transfected with shSertad1-zsgreen and shRand-zsgreen (scrambled RNA) and treated with 1.5 µM Aβ for the indicated time points 48 hours post transfection. The number of live green neurons were counted under fluorescence microscope. Representative image of one such neuron transfected with shSertad1 and shRand is depicted upto 48 h of Aβ treatment. (**B**) Graphical representation of percent (%) viable cells. Asterisks denote statistically significant differences between shSertad1 and shRand transfected cells. Data represent Mean±SEM of five independent experiments. (**C**) Tissue slices (30 µm) were prepared from brain expressing shSertad1 and EV expressing lentiviral particles. TUNEL assay was performed 21 days post-surgery as per manufacturer’s protocol. Representative image showing TUNEL positive cells (red puncta). Nuclei were stained with Hoechst (blue) (magnification: 63X). (**D**) Graphical representation of % TUNEL positive cells (N=3 animals/group). Tissue slices from brains expressing shSertad1 or EV in 5xFAD or WT mice were stained with 6E10 antibody. (**E**) Left panel shows 6E10 stained image and right panel shows merged image. Images were taken at 63X magnification. (**F**) Graphical representation of number of plaques [**F(i)]**, Aβ plaque burden [**F(ii)**] and Amyloid integrated density [**F(iii)**]. Data represent Mean±SEM. Statistical analysis was done using One-way ANOVA, Tukey’s post-hoc analysis. Asterisks denote statistically significant differences between indicated groups; *p<0.05, **p<0.01 and ***p<0.001.

### Sertad1 plays a crucial role in dysfunctional autophagy observed in AD transgenic mice

There are very few studies that suggest a link between Sertad1 and autophagy-lysosomal pathway in cancer [22, 24]. Since Sertad1 promotes apoptosis and there exists a complex yet unexplored cross-talk between these two cell death modalities, we hypothesized that it might have a role in autophagy modulation. We sacrificed the shSertad1 or EV injected 5xFAD and WT mice after 21 days post-surgery. We isolated the cortex and hippocampus from these mice and subjected them to biochemical analysis. We found that LC3II:LC3-I ratio and LC3II: actin, that was increased in case of 5xFAD mice, was significantly reduced upon Sertad1 knockdown in both hippocampal and cortical tissue lysates (Figure 5A, 5B and 5C) suggesting a reduction in autophagosome number. We assessed p62 levels, and found that it was elevated in 5xFAD mice, and significantly reduced upon Sertad1 knockdown in hippocampus and cortex of 5xFAD mice (Figure 5A and 5D). Next, we checked ATG5 levels, which is a marker for autophagosome elongation and found that it was elevated in 5xFAD mice and significantly reduced upon Sertad1 interference (Figure 5A and 5E). Furthermore, we assessed LAMP1 protein levels, lysosomal membrane marker and found that it was upregulated in 5xFAD mice and was significantly lowered upon Sertad1 knockdown in both hippocampal and cortical tissue lysates (Figure 5A and 5F). This is in agreement with our results obtained from Aβ treated primed PC12 cells where we transfected shSertad1-zsgreen and shRand-zsgreen and treated them with oligomeric Aβ followed by immunocytochemical analysis to check LC3 and p62 expression. We found that shSertad1 transfected cells show low expression levels of LC3 and p62 in immunofluorescence studies and reduced number of puncta of LC3 and p62 respectively as compared to scrambled RNA transfected cells (Fig. S1A-1H). This suggests that Sertad1 knockdown reduces number of accumulated autophagosomes possibly by enhancing autophagy flux.

**Figure 5:**
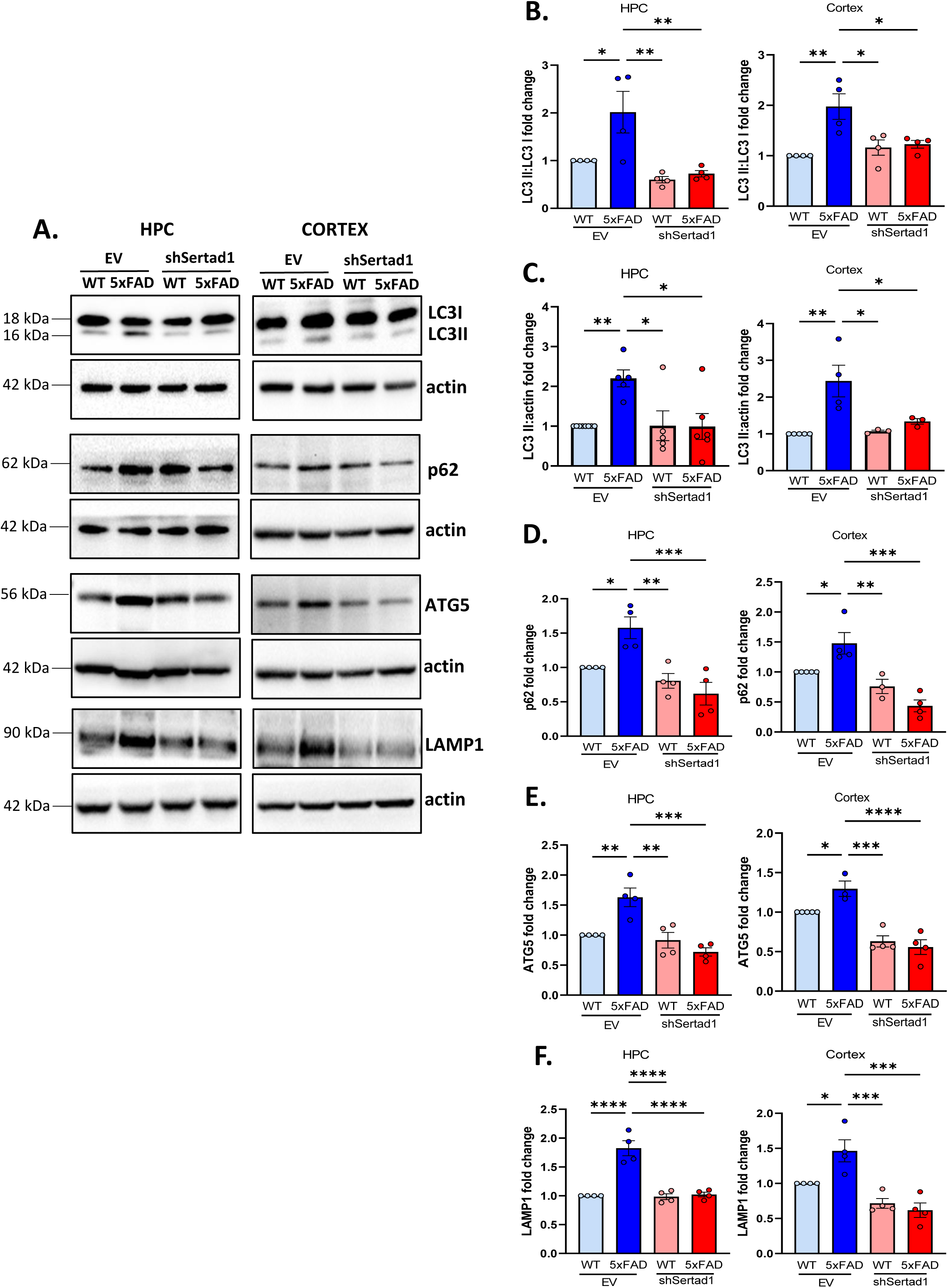
Sertad1 knockdown ameliorates autophagy dysfunction in in-vitro and in-vivo models of AD. shSertad1 and EV expressing lentiviral particles were stereotactically infused in 5xFAD and WT mice brain, protein lysates from cortex and hippocampus were prepared 21 days post-surgery and subjected to Western blotting. (**A**) Representative immunoblots showing expression levels of autophagosome markers: LC3, p62 and ATG5, and lysosomal marker: LAMP1 in hippocampal and cortical lysates of shSertad1 or EV infused 5xFAD and WT mice 21 days post-surgery as indicated. Actin was used as loading control. Graphical representation of fold change of LC3II:LC3I (**B**), LC3II (**C**), p62 (**D**), ATG5 (**E**) and LAMP1 (**F**). (N=4 animals/group). Data represent Mean±SEM. Statistical analysis was done using One-way ANOVA, Tukey’s post-hoc analysis. Asterisks denote statistically significant differences between indicated groups; *p<0.05, **p<0.01, ***p<0.001 and ****p<0.0001.

### Sertad1 knockdown blocks FoxO3a translocation to the nucleus and restores Akt activity that is lost in AD pathogenesis

We next explored the mechanism by which Sertad1 modulates autophagy mediated cell death in AD. FoxO3a has been reported to directly regulate autophagy genes as seen in adult neural stem cells by Chip-seq through transcriptional regulation [41]. Apart from transactivation functions, FoxO3 is known to activate FoxO1 dependent autophagy, thus playing an indirect role in autophagy modulation [42]. Cytosolic FoxOs can regulate autophagy by binding to autophagy proteins like ATG7 and epigenetically through histone modifications and microRNAs [43]. We have reported that FoxO3a is dephosphorylated at the Akt phosphorylation site: Ser 253 and translocates to the nucleus upon Aβ treatment [44]. FoxO3a translocation and activation is finely controlled through dynamic involvement of 14-3-3 and Akt by protein phosphatase 2A [45]. We asked the question whether Sertad1 is required for FoxO3a localization and activation. To assess FoxO3a localization, we transfected primed PC12 cells with shSertad1-zsgreen and shRand-zsgreen, maintained the cells for 48 hours followed by 3 µM Aβ treatment and then immunostained with FoxO3a. Our results revealed that in shSertad1-zsgreen transfected cells, FoxO3a was retained exclusively in the cytosol as compared to shRand-zsgreen transfected cells, where FoxO3a translocated to the nucleus in response to Aβ exposure (Figure 6A-6C). We next assessed phosphorylation of FoxO3a at Ser253 and found reduced phosphorylation in 5xFAD mice which was restored significantly in shSertad1-EGFP infused 5xFAD mice in both hippocampal and cortical lysates (Figure 6D and 6E). Interestingly, our study revealed that Sertad1 knockdown not only blocked FoxO3a nuclear translocation but also reduced total protein levels of FoxO3a (Figure 6D and 6F). It is known that Akt, a survival kinase, remains inactive in the AD brain as indicated by reduced phosphorylation [46]. Akt is an upstream negative regulator of FoxO3a [45]. We have observed reduced Akt phosphorylation in 5xFAD mice as compared to shSertad1-eGFP injected 5xFAD mice where Akt phosphorylation at Thr 308 is significantly restored both in cortical and hippocampal lysates (Figure 6G and 6H). Our results show that Sertad1 modulates autophagy through the Akt/FoxO3a axis and Sertad1 knockdown not only inhibits FoxO3a translocation to the nucleus but also reduces its protein levels. Further, Akt, a survival kinase whose activity is lost in 5xFAD mice is restored in shSertad1 infused mice.

**Figure 6:**
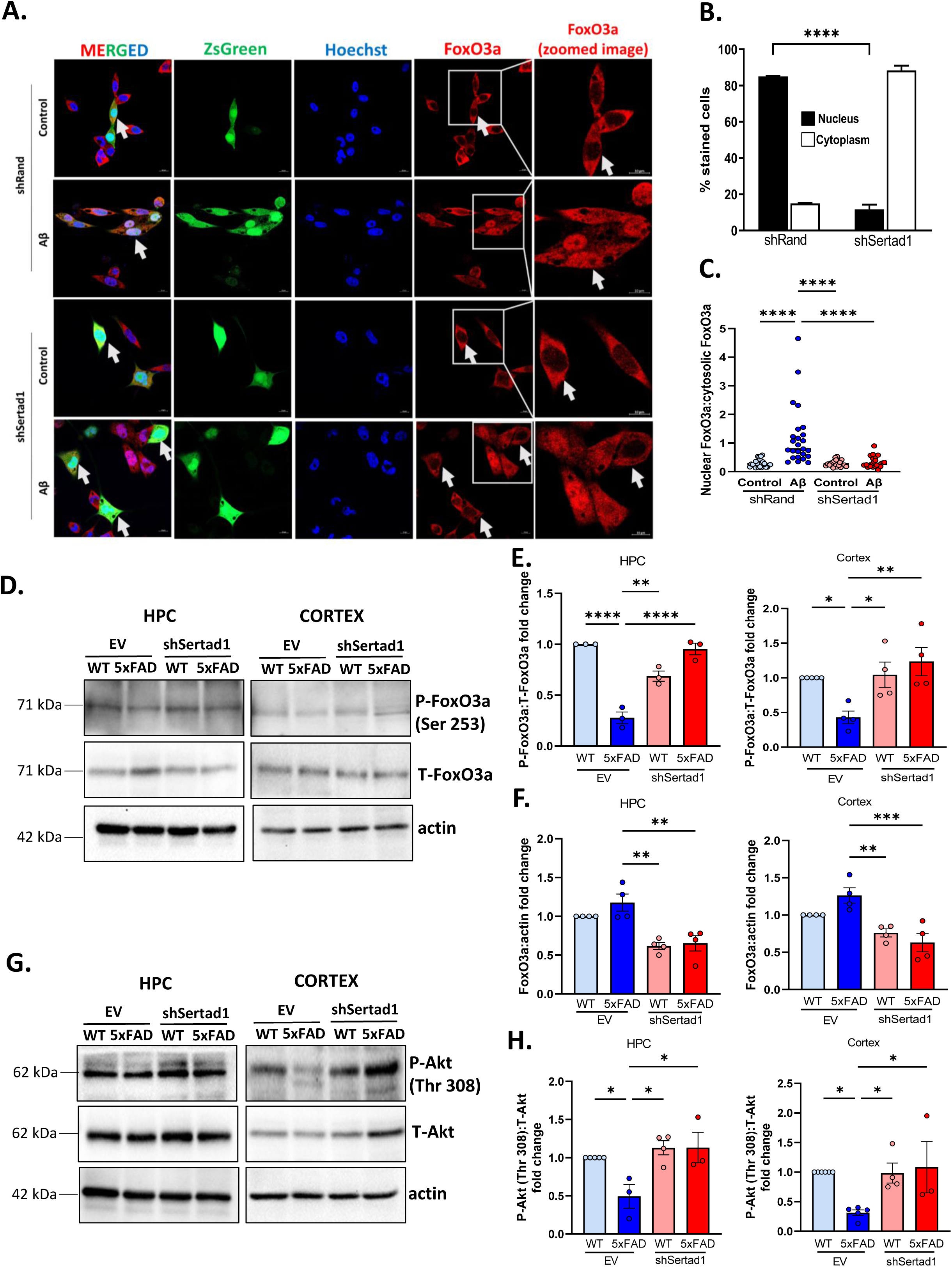
Sertad1 aids in FoxO3a translocation to the nucleus and subsequent activation of autophagy genes by inhibiting Akt activity. (**A**) PC12 cells were transfected with pSIREN-shSertad1-zsgreen (shSertad1) and pSIREN-shRand-zsgreen (shRand), primed and then treated with 3 µM Aβ for 16 hours. Representative images of transfected PC12 cells treated with Aβ along with respective controls. Left to right: first panel shows immunostaining of FoxO3a (red), second panel shows zsgreen (green), third panel shows nuclei stained with Hoechst (blue), fourth panel shows merged image and fifth panel shows enlarged image of FoxO3a staining (magnification 63X). (**B**) Percentage-stained cells indicate the proportion of transfected cells where FoxO3a is present in the cytosol and proportion of transfected cells where FoxO3a is present in the nucleus. Graphical representation of % stained cells are depicted. Approximately 50 cells were counted per experiment. (**C**) Graphical representation of nuclear FoxO3a: cytosolic FoxO3a. (**D**) Representative immunoblots showing P-FoxO3a and FoxO3a protein levels in hippocampus and cortex of shSertad1 or EV infused 5xFAD mice and WT mice 21 days post stereotaxic surgery. Actin was used as loading control. Graphical representations of fold change of P-FoxO3a: T-FoxO3a (**E**) and fold change of FoxO3a with respect to actin (**F**) in hippocampus and cortex of mice of indicated groups analysed by densitometry using NIH-ImageJ. (N=3-4 animals/group). (**G**) Representative immunoblots showing fold change of P-Akt (Thr308):T-Akt and actin in hippocampal and cortical lysates of shSertad1 or EV infused 5xFAD and WT mice 21 days post-surgery. (**H**) Graphical representations of fold change of P-Akt:T-Akt in hippocampus and cortex analysed by densitometric analysis using NIH-ImageJ software. (N=4 animals/group). Data represent Mean±SEM. Statistical analysis was done using One-way ANOVA, Tukey’s post-hoc analysis. Asterisks denote statistically significant differences between indicated groups; *p<0.05, **p<0.01, ***p<0.001 and ****p<0.0001.

### Sertad1 knockdown ameliorates cognitive deficits that are observed in the 5xFAD mice brain

Our next aim was to assess whether Sertad1 knockdown in 5xFAD mice can reduce the behavioural deficits that are observed in AD transgenic mice. A battery of behavioural experiments was performed 14 days post-injection of shSertad1-eGFP and EV-eGFP lentiviral particles in the CA1 region of hippocampus of 5xFAD and WT mice. Mice were grouped as (1) EV-WT, (2) EV-5xFAD, (3) shSertad1-WT and (4) shSertad1-5xFAD. First, we performed the open field test or locomotion test where we assessed the locomotor activity and thigmotaxis behaviour. Degree of thigmotaxis is considered a measure of anxiety and WT animals tend to remain close to the walls of the open field. Our results indicate that 5xFAD mice show hyper-locomotor activity and reduced thigmotaxis behaviour as evident from the infrared recording images, total distance travelled in cm and time spent in outer zone in seconds, which is attenuated in shSertad1-eGFP injected 5xFAD mice when given a total exploration time of 10 minutes (Figure 7A-C). Figure 7D and 7F show schema for contextual and cue dependent fear conditioning on the training day respectively. In both the experiments, we calculated the average % freezing where mice received foot-shock repeated for 4 test sessions on the acquisition day. Details of the protocol are given in the materials and methods section. No foot-shock was given on the test day. Mice which retain memory of receiving a foot-shock, freeze in anticipation of receiving a shock, even when there is no shock given on the probe day, showing higher % freezing. In the contextual fear conditioning test, we observed that 5xFAD mice show significantly lower % freezing due to lower memory retention as compared to shSertad1-eGFP 5xFAD mice, where % freezing was significantly restored (Figure 7D and 7E). Similarly, in the cue dependent fear conditioning test, we observed that 5xFAD mice show less % freezing, which is significantly restored in shSertad-eGFP injected 5xFAD mice, indicating Sertad1 knockdown restores cue dependent fear-based memory in 5xFAD mice (Figure 7F and 7G). Our results indicate that regions of the brain associated with fear based associative learning, that is, amygdala and hippocampal coordination is significantly improved in shSertad1 injected 5xFAD mice as compared to EV injected 5xFAD mice. Next, we performed the Novel Object Recognition (NOR) task that assesses long term recognition memory. Figure 7H shows the schema for the experiment on the test day (Day 2). We exposed the mice to an open arena and allowed them to explore two similar shaped objects and recorded the time spent near each object, when given a total exploration time of 5 minutes (Figure 7I). On the next day, we replaced one of the objects with a differently shaped novel object and recorded the time spent by mice near each of the objects given a total time of 5 minutes. Our results show that EV injected 5xFAD mice are unable to differentiate between novel and familiar objects, and thus show no significant difference between time spent near novel and familiar objects as also evident from negative discrimination index values. On the contrary, shSertad1-eGFP injected 5xFAD mice spend significantly more time near the novel object (Figure 7J) and display a positive value for discrimination index indicating that Sertad1 knockdown restores long term recognition memory in 5xFAD mice (Figure 7K). We next performed the Morris Water Maze test which is a classic experiment to test hippocampus dependent spatial navigation and memory [47]. We trained the animals to find the hidden platform for 3 days, where four trials were given to each animal per day (a total of 12 trials were given) and recorded the average escape latency of four trials on day 1, day 2 and day 3 in seconds. Details of experimental setup is given in the materials and methods section. Figure 7L depicts a pictorial representation of the path traversed by mice on the probe day of one mouse from each group. Our results clearly indicate that 5xFAD mice take longer time to find the hidden platform as compared to shSertad1-eGFP injected 5xFAD mice which take significantly lesser time to find the platform on the second and third days (Figure 7M). On the fourth day, the hidden platform was removed and mice were allowed to explore the maze for 90 seconds. Time spent by mice in the target quadrant and number of target quadrant entries was recorded. We observed that while 5xFAD mice swim across all the quadrants, shSertad1-eGFP injected mice spend more time where they anticipate the platform to be located. EV injected 5xFAD mice spend less time in the target quadrant and show less number of target quadrant entries as compared to shSertad1-eGFP injected mice which spent significantly higher time in the target quadrant and entered the target quadrant a greater number of times in anticipation of finding the platform on the probe day (Figure 7N and 7O). This data clearly shows that Sertad1 knockdown ameliorates various aspects of impaired cognitive functions including anxiety like behaviour, long term recognition memory, fear based associative learning, spatial and navigation learning that are observed in 5xFAD mice. These results demonstrate a strong link between amelioration of autophagy deficits and improvement in cognitive behaviour in 5xFAD mice

**Figure 7:**
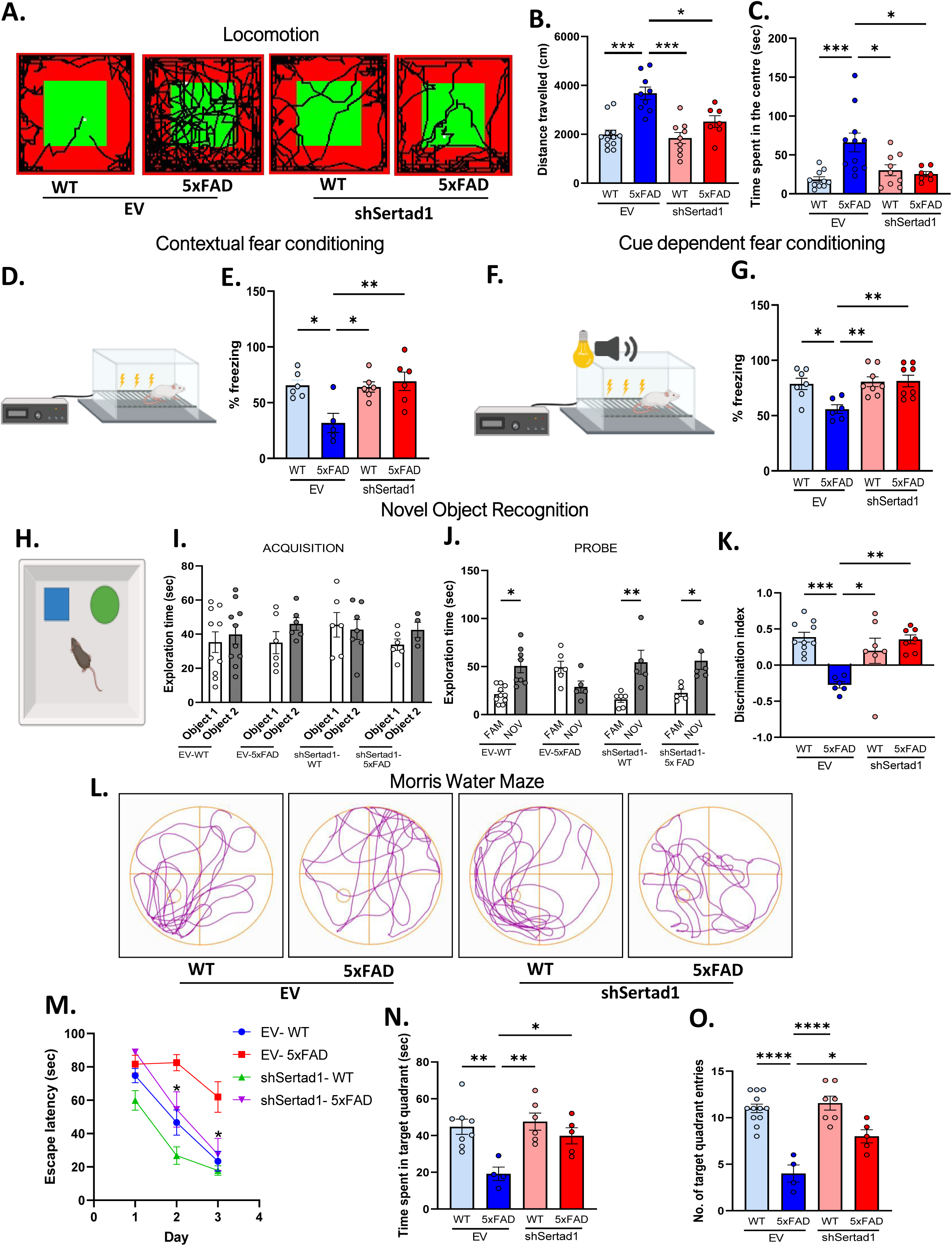
Sertad1 knockdown ameliorates behavioural deficits in 5xFAD mice. Battery of behavioural experiments were performed with animals divided into four groups: (1) EV expressing lentiviral particles infused in WT mice (EV-WT), (2) EV expressing lentiviral particles infused in 5xFAD mice (EV-5xFAD), (3) shSertad1 expressing lentiviral particles infused in WT mice (shSertad1-WT) and (4) shSertad1 expressing lentiviral particles infused in 5xFAD (shSertad1-5xFAD) mice. Behavioural experiments were performed 14 days post-surgery. Number of animals used for behavioural experiments in each group is given in tabular form in the supplementary section. (**A**) Representative infrared images of path traversed by a mouse in each group in the Open Field Test. (**B**) Graphical representation of average total distance travelled by each mouse in cm. (**C**) Graphical representation of time spent in the centre by animals of indicated groups in seconds. (**D**) Representation of experimental setup for contextual fear conditioning experiment. (**E**) Graphical representation of percentage freezing of mice in indicated groups recorded one day after training. (**F**) Representation of experimental setup for cue dependent fear conditioning experiment. (**G**) Graphical representation of percentage freezing of mice in indicated groups recorded one day after training. (**H**) Diagrammatic representation of experimental setup for Novel Object Recognition (NOR). (**I**) Graphical representation of time spent near objects in seconds by mice of indicated groups on the acquisition day when given 5 minutes of total exploration time. (**J**) Graphical representation of time spent near novel and familiar objects respectively by mice of indicated groups on the probe day when given 5 minutes of total exploration time. (**K**) Discrimination index of mice is calculated as follows: Discrimination index = (time spent near novel object-time spent near familiar object)/ (time spent near novel object+ time spent near familiar object). Graphical representation of discrimination index of mice of indicated groups. (**L**) Diagrammatic representation of path traced by each mouse of the indicated groups using ANY-MAZE software in the Morris Water Maze experiment on the probe (fourth) day. (**M**) Graphical representation of escape latency indicating time taken in seconds to find the hidden platform at the indicated time points. (**N**) Graphical representation of time spent in target quadrant (in seconds) of mice of indicated groups when given a total exploration time of 90 seconds on the probe (fourth) day in the water maze. (**O**) Graphical representation of number of target quadrant entries of mice of indicated groups when given a total exploration time of 90 seconds on the probe (fourth) day in the water maze. Data represent Mean±SEM. Statistical analysis was done using one-way ANOVA, Tukey’s post-hoc analysis. Asterisks denote statistically significant differences between indicated groups; *p<0.05, **p<0.01, ***p<0.001 and ****p<0.0001.

### Sertad1 knockdown ameliorates synaptic deficits and restores dendritic spine density in 5xFAD mice

Since absence of Sertad1 ameliorated cognitive deficits as evident from different behavioural parameters that we tested in 5xFAD mice, we were interested to explore the biochemical basis of this recovery. Learning and memory involves the formation of new dendritic spines and considerable alternations in spine dynamics [48]. Synaptic dysfunction is among the earliest events in 5xFAD mice brain pathogenesis. Li et al. reported that the decline in hippocampal LTP occurs at 10-12 weeks of age in 5xFAD mice. Different aspects of learning and memory deficiencies in 5xFAD mice become apparent at 4-5 months of age [13]. Thus, we were interested to assess the expression levels of pre- and post-synaptic proteins to see if Sertad1 knockdown has any effect on protein levels of synaptic proteins. PSD95 or postsynaptic density 95, is a post-synaptic scaffolding protein localized primarily in excitatory synapses and loss in PSD95 levels is associated with neurodegeneration [49]. SNAP25 or synaptosomal-associated protein of 25 kDa is a pre-synaptic protein that helps in organization of the molecular apparatus that promotes spine formation [50]. We sacrificed mice 21 days after infusion of shSertad1-eGFP and EV-eGFP and checked for pre- and post-synaptic proteins. As expected, we found that PSD95 and SNAP25 protein levels were low in 5xFAD mice, but was significantly restored in both hippocampal and cortical whole cell lysates of 5xFAD mice injected with shSertad1-eGFP (Figure 8A-D). The size, morphology and density of dendritic spines that constitute majority of excitatory synapses strongly correlates with cognitive functions [51]. We assessed the cytoarchitecture of the brain by Golgi-cox staining to quantify dendritic spine density. Detailed procedure for Golgi-cox staining is given in the materials and methods section. We perfused mice 21 days after shSertad1-eGFP and EV-eGFP injection in 5xFAD mice brain and isolated the brains. Figure 8E depicts an image of the brain area that shows the cytoarchitecture of dendritic spines of cortex and hippocampus. The density and morphology of the dendritic spines of cortex and hippocampus are shown in figure 8F in an enlarged image. Concomitant to the restoration in pre- and post-synaptic proteins upon Sertad1 knockdown, we found that shSertad-eGFP injected mice show a significant restoration in spine number quantified per 10 µm length as compared to 5xFAD mice injected with EV-eGFP where spine density is low (Figure 8F and 8G). Thus, our results reveal that Sertad1 knockdown restores pre- and post-synaptic protein levels, PSD95 and SNAP25 respectively and dendritic spine density in 5xFAD mice.

**Figure 8:**
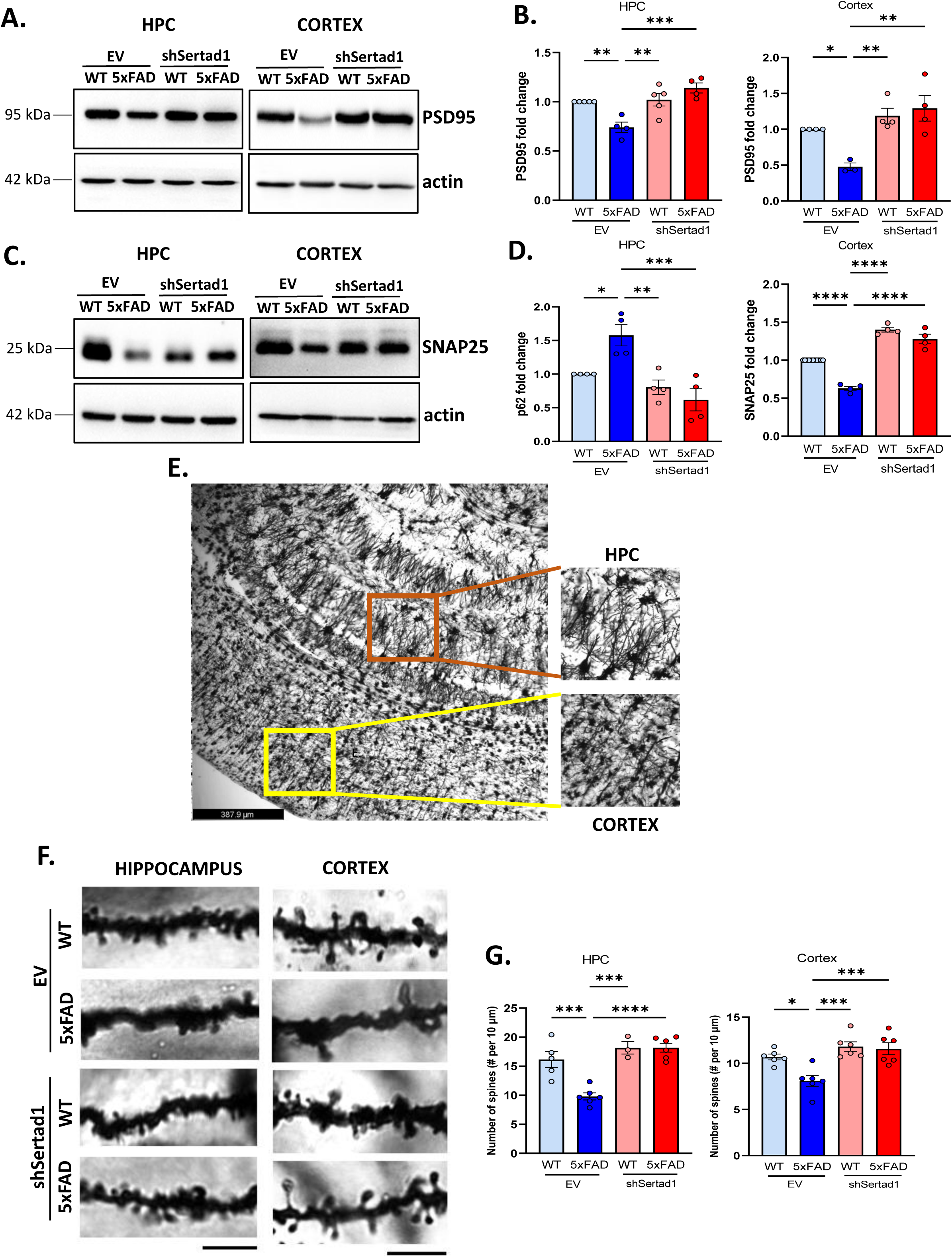
Sertad1 knockdown restores synaptic deficits and dendritic spine density in 5xFAD mice. (**A**) Representative immunoblots of PSD95 expression levels in hippocampal and cortical lysates of shSertad1 or EV infused 5xFAD and WT mice as indicated, 21 days post-surgery. (**B**) Graphical representation of PSD95 fold change in hippocampus and cortex is depicted. Actin was used as loading control. (**C**) Representative immunoblots of SNAP25 expression levels in hippocampal and cortical lysates of shSertad1 or EV infused 5xFAD and WT mice 21 days post-surgery. (**D**) Graphical representation of SNAP25 fold change in hippocampus and cortex is depicted. Actin was used as loading control. (**E**) Representative image of Golgi-cox staining brain slices (75 µm) prepared from shSertad1 or EV infused 5xFAD and WT mice 21 days post-surgery at 10X magnification. Right panel shows zoomed image of cortex and hippocampus as indicated. (**F**) Representative image of dendritic spines in hippocampus and cortex of tissue slices (75 µm) of shSertad1 or EV infused 5xFAD and WT mice 21 days post-surgery in the hippocampus and cortex. Scale bar=5 µm. (**G**) Graphical representation of dendritic spine number per 10 µm dendritic length in the hippocampus and cortex of shSertad1 or EV infused 5xFAD and WT tissue sections. Data represent Mean±SEM. Statistical analysis was done using one-way ANOVA, Tukey’s post-hoc analysis. Asterisks denote statistically significant differences between indicated groups; *p<0.05, **p<0.01, ***p<0.001 and ****p<0.0001

## Discussion

In this study, we assessed autophagy progression with age by analysing protein level of markers of initiation, elongation and maturation of autophagosomes and lysosomes in 5xFAD mice and found their robust induction. Proponents of the amyloid cascade hypothesis, first proposed by Hardy and Higgins consider that Aβ deposition is the first event that triggers further pathogenic alterations in signaling pathways downstream of plaque deposition [2]. However, considering long incubation time of the disease pathogenesis and clinical failures in targeting Aβ directly led to a speculation that proteopathic stress exerted by Aβ aggregation and neurofibrillary tangles disrupts the homeostasis of the brain by altering normal cellular functions leading to clinical manifestation of dementia [8, 9]. A comprehensive age dependent study of 5xFAD mice has shown that hippocampal LTP decline and synaptic impairments begin at 10 weeks of age in 5xFAD mice when plaque deposition has not yet begun [13]. An important question is whether age related deficiencies in protein aggregate clearance mechanisms results in their neurotoxic accumulation or aggregate formation overwhelms the proteostatic system to hinder the clearance process [8]. It has been reported that autophagy is impaired during the later stages, and a failure in lysosomal acidification results in an inability to clear up cellular waste [33, 52]. Increased synthesis of autophagosomes and lysosomes in response to aggregate stress, overburdens failing lysosomes with deficient lysosomal proteases resulting in neurite dystrophy [34, 53] It is difficult to delineate specifically the causal relationship between the pathogenic events in AD. These cellular and molecular changes influence each other. Overwhelming of homeostatic strategies of the cell results in the manifestation of cognitive decline and irreversible neuronal loss in the brain that neurons are unable to cope up with. Failure to maintain proper balance between protein synthesis and aggregate clearance mechanisms probably lead to the onset of AD pathogenesis.

Autophagy and apoptosis have some common inducers and the two processes are involved in an intricate cross-talk with each other [18, 19]. Our previous studies have revealed simultaneous induction of autophagy and apoptosis with AD progression but the precise mechanism of this cross-talk is not well known. We have identified an endogenous protein, Sertad1 that is known to play a role in various cellular processes such as cell cycle modulation, transcriptional regulation, senescence inhibition and cancer progression [24, 25, 54–57]. Sertad1 is primarily a transcriptional coregulator of E2F1 that binds and activates CDK4, thereby activating the apoptotic pathway by phosphorylation of Rb resulting in dissociation and derepression of a protein complex comprising of transcription factor proteins and chromatin modifiers and activation of Myb and Bim, downstream targets of E2F1 [20]. There are studies that suggest an association of Sertad1 with cell death paradigms like autophagy, apoptosis, and necrosis in cancer models [22, 24, 58–60]. We have addressed if Sertad1 has an important role in dysfunctional autophagy in AD mice and established its role and molecular mechanism in dysfunctional autophagy. Several studies have established FoxO3a as a master regulator of the autophagy network through direct as well as indirect means and acts as a link between autophagy and apoptosis pathways [41, 42, 61]. The subcellular localization of FoxO3a is finely controlled through dynamic and reversible phosphorylation and dephosphorylation events controlled by specific kinases and phosphatases [44, 45]. Under normal conditions, FoxO3a is phosphorylated at Ser 253, that overlaps with the nuclear localization signal (NLS) impeding its import into the nucleus. In AD, however, FoxO3a is dephosphorylated and translocates to the nucleus to activate autophagy and apoptosis genes [44]. Interestingly, we found that shSertad1 injected mice showed a restoration in phosphorylation of FoxO3a at Ser 253, thus causing its cytosolic retention, but also showed a reduction in protein levels. As previously discussed, Sertad1 binds to E2F1, thus regulating its transcriptional activity. FoxO transcription factors and E2F1 function in a feed-forward loop that regulate certain common target genes [62]. Further, E2F1 regulates FoxO protein expression by binding to its promoter site [63]. Akt is a survival kinase that negatively regulates FoxO3a activity as well as E2F1 dependent apoptotic program. We found that Akt activity is also restored upon Sertad1 knockdown. Sertad1 is known to interact with protein phosphatases like PP2A [29]. A possible mechanism by which Sertad1 regulates the phosphorylation state of Akt and FoxO3a could be through binding to and regulating the phosphatase activity of PP2A. Thus, we can say that Sertad1 is a multimeric protein that interacts with different mediators and post-translational modulators lying upstream of this regulation, thus playing an important role in the decisive process of cell death.

Further, Sertad1 inactivation restores learning and memory deficits in 5xFAD mice, where we found a remarkable rescue in fear based cognitive learning and spatial learning tasks along with a restoration in synapse health upon its knockdown. Learning results in a surge in protein synthesis which needs to be balanced through degradation mechanisms like autophagy. Autophagy is crucial for long term memory consolidation and dendritic spine plasticity [64]. Autophagy is reduced during aging and its stimulation in hippocampal neurons restores cognitive loss associated with aging [65]. Another recent study has shown that autophagy deactivation leads to loss of synaptic homeostasis with implications in AD pathogenesis [66]. Given the importance of autophagy mediated degradation mechanisms in cognition, it is puzzling to understand how its dysfunction can lead to pathogenic changes in the AD brain. Our results open new avenues to explore how autophagy acts as a double-edged sword in the context of cognitive learning and synapse biology. Neurons being highly compartmentalized structures adds another layer of complexity to region specific regulatory mechanisms. Furthermore, since the brain has different types of cells, including microglia, astrocytes and oligodendrocytes, it will be interesting to explore how different signalling mechanisms interact with each other.

Nonetheless, our results clearly prove the neuroprotective potential of transcriptional coregulator, Sertad1. This protein is a master regulator that lies upstream of many crucial signalling pathways that induce cell death. Among its multifaceted roles, we have established its regulatory function in dysfunctional autophagy in 5xFAD mice. We have also shown its role in improvement in learning and memory functions through the stable delivery of lentiviral particles expressing shSertad1 that opens new avenues for further exploration. This study, thus highlights that Sertad1 is an excellent target for therapeutic intervention for ameliorating AD pathology.

## Conclusion

We show that Sertad1 is elevated in 5xFAD mice brain, which is concomitant with aberrant autophagy induction that worsens as the disease progresses (Figure 1). AD is a multifactorial disorder with several pathways going awry simultaneously resulting in complete loss of homeostasis [8]. Since neurons rely so heavily on autophagy for protein aggregate clearance, it is imperative to understand the exact mechanism of autophagy flux dysfunction. We propose a model where Sertad1, a proapoptotic molecule regulates aberrant autophagy in 5xFAD mice through the Akt/FoxO3a axis and its downregulation by delivering lentiviral particles expressing shSertad1 can rescue deficits in cognition along with restoration in dendritic spine density and synapse health in 5xFAD mice (Figure 9). Thus, our study brings to light, an excellent target for therapeutic intervention to combat debilitating disorders like AD by use of a stable genetic manipulation approach.

**Figure 9:**
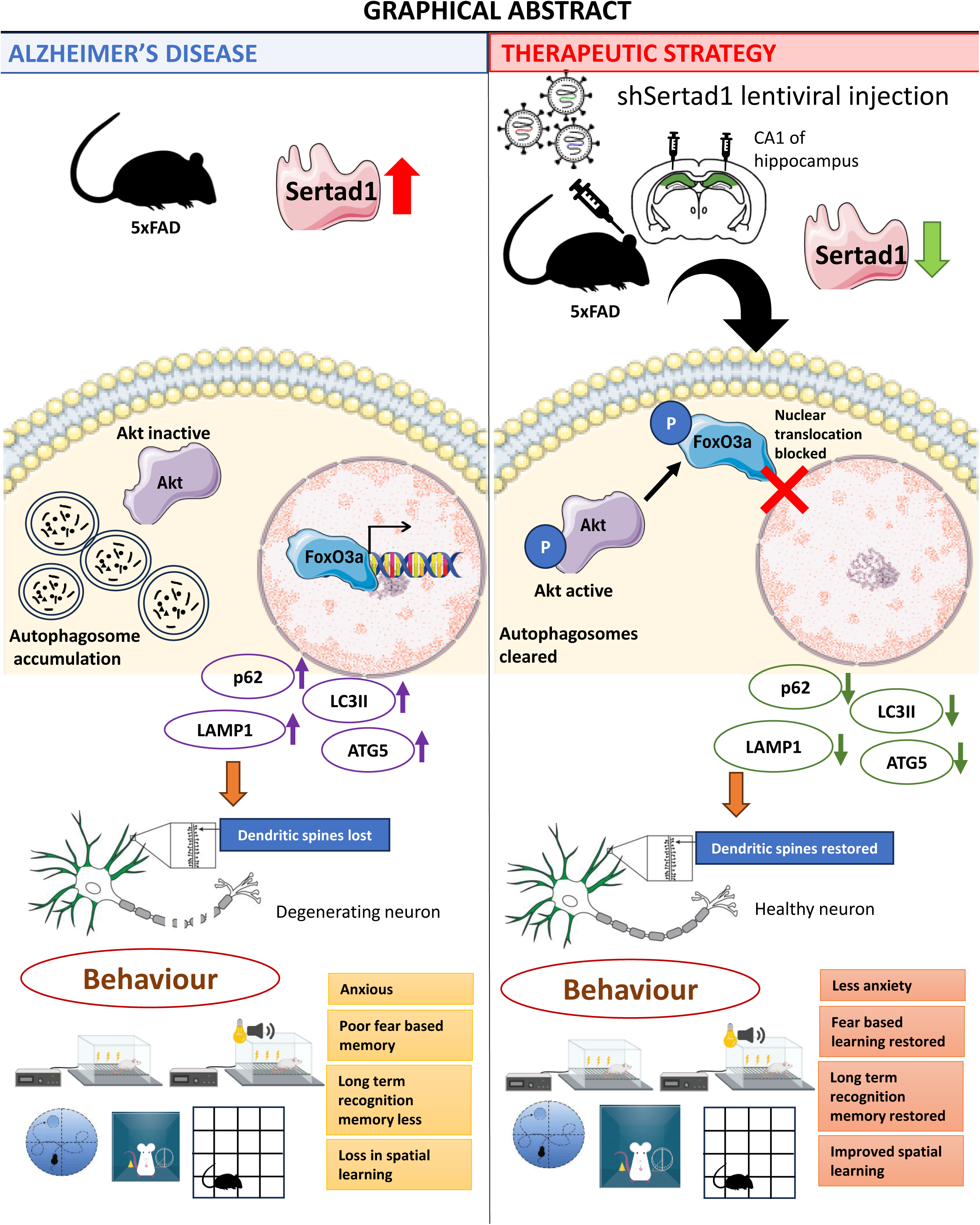
Schematic representation depicting that Sertad1 interference using RNAi can ameliorate cognitive and synaptic deficiencies in 5xFAD mice brain. Graphical abstract shows that autophagosome synthesis is robustly induced in 5xFAD mice. Sertad1 is upregulated in 5xFAD mice in an age-dependent manner. Knockdown of Sertad1 ameliorates synaptic dysfunction, neurodegeneration and cognitive deficiencies in the 5xFAD mice.

## METHODS

### Materials

Dulbecco’s modified Eagle’s medium (DMEM)-F12, DMEM high glucose medium, Neurobasal medium, OPTI-MEM medium, B27 supplement, Fetal Bovine Serum (FBS), Horse serum (HS), Normal goat serum (NGS) Penstrep antibiotic, Lipofectamine 2000, Hoechst 33342, and Prolong Gold antifade with DAPI were purchased from Thermo Fisher Scientific (MA, USA). Human recombinant Nerve Growth Factor (NGF), insulin, progesterone, putrescine, selenium, apo-transferrin, poly-D-lysine (PDL), radioimmunoprecipitation assay (RIPA buffer), 1,1,1,3,3,3 hexafluoro-2-propanol (HFIP), paraformaldehyde (PFA), dimethyl sulphoxide (DMSO), PhosSTOP and bovine serum albumin (BSA) were purchased from Sigma (St. Louis, MO, USA). Protease inhibitor cocktail, Lenti-X concentrator, enhanced chemiluminescence (ECL), Western blot stripping buffer and protein ladder were purchased from Takara bio (Kusatsu, Japan). 0.45 µm Polyvinylidene (PVDF) membrane was purchased from Merck Millipore (Darmstadt, Germany). Other fine chemicals and reagents were procured from standard local suppliers. List of antibodies that have been used in this work along with their dilution and application is given in the table below:

**Table.** List of antibodies. WB = western blot, ICC = immunocytochemistry, IHC = immunohistochemistry, CST = Cell Signalling Technology, SCBT = SantaCruz Biotechnology

| **Antibody** | **Species**<br>**raised in** | **Application** | **Dilution** | **Source** | **Catalogue**<br>**number** |
| --- | --- | --- | --- | --- | --- |
| phospho-Akt<br>(Thr 308) | rabbit | WB | 1:500 | CST | 4056S |
| Akt | rabbit | WB | 1:1000 | CST | 9272S |
| 6E10-APP | rabbit | IHC | 1:1000 | Novus | NBP2-62566 |
| Phospho-FoxO3a<br>(Ser253) | rabbit | WB | 1:500 | CST | 9466S |
| FoxO3a | rabbit | WB<br>ICC | 1:1000<br>1:100 | CST | 2497S |
| Sertad1 | rabbit | WB<br>IHC | 1:500<br>1:100 | Invitrogen<br>Novus | PA5-75674<br>NBP1-31052 |
| LC3B | rabbit | WB<br>ICC | 1:1000<br>1:100 | Novus | NB100-2220 |
| p62 | rabbit | WB<br>ICC | 1:1000<br>1:100 | R&Dsystems | MAB8028 |
| LAMP-1 | rabbit | WB | 1:1000 | Abcam | ab24170 |
| ATG5 | rabbit | WB | 1:1000 | Novus | NB110-53818 |
| PSD95 | rabbit | WB | 1:2000 | Abcam | ab18258 |
| SNAP25 | mouse | WB | 1:1000 | SCBT | sc20038 |
| GFP | mouse | WB<br>IHC | 1:1000<br>1:200 | SCBT | sc9996 |
| Anti-rabbit<br>IgG-HRP<br>linked<br>antibody | goat | WB | 1:5000 | CST | 7074S |
| Anti-mouse<br>IgG-HRP<br>linked<br>antibody | horse | WB | 1:5000 | CST | 7076S |

### Cell culture

Rat pheochromocytoma (PC12) cells were cultured in DMEM high glucose supplemented with 10% heat in-activated horse serum (HI-HS) and 5% heat-inactivated fetal bovine serum (HI-FBS) as described previously [67]. Naïve PC12 cells were differentiated using NGF at 100 ng/ml concentration with 1% HI-HS for three days prior to treatment. Human embryonic kidney 293T (HEK293T) cells were cultured in DMEM high glucose medium supplemented with 10% HI-FBS. Primary rat cortical neurons were cultured as described previously [68, 69]. In brief, cells from the neocortex of E18 days pregnant Sprague Dawley rat foetus were dissected under a dissection microscope. Isolated cells were subjected to gentle trituration and then plated on poly-D-lysine coated plates. Cultured cells were maintained in-vitro in DMEM-F12 medium supplemented with D-glucose (6mg/ml) and apo-transferrin (0.1 mg/ml), insulin (25 µg/ml), progesterone (20 ng/ml), putrescine (60 µg/ml) and selenium (30 ng/ml). After maintaining cells for 24 hours, half of the cortical medium was replaced with Neurobasal medium supplemented with B27 supplement. Cortical neurons were treated with 1.5 µM Aβ on the 6^th^ day of plating in fresh Neurobasal medium containing B27.

### Preparation of oligomeric Amyloid-β (Aβ)

Lyophilized synthetic Aβ (1-42) peptide with sequence: DAEFRHDSGYEVHHQKLVFFAEDVGSNKGAIIGLMVGGVVIA (Cat no. AB-250-10) was purchased from AlexoTech (Vasterbottens Ian, Sweden). Lyophilized peptide was dissolved in 100% HFIP solvent to a final concentration of 1 mM and centrifuged in a speed-vac (Eppendorf, Hamburg, Germany). Peptide pellet was then resuspended in DMSO to a final concentration of 5 mM followed by sonication at 37°C for 10 minutes. Obtained solution was then stored in small aliquots at -80°C. Prior to use, aliquots were diluted in phosphate buffered saline (PBS: 137 mM NaCl, 2.7 mM KCl, 10 mM Na_2_HPO_4_, 2 mM KH_2_PO_4_, pH 7.4) and 0.2% sodium dodecyl sulphate (SDS) to a final concentration of 400 µM and incubated at 37°C for 6-18 hours. The solution was further diluted with PBS to a final concentration of 100 µM and incubated at 37°C for 18-24 hours.

### Aβ treatment on cells

Naïve PC12 cells were treated with 3 µM Aβ (1-42) after priming for 3 days for indicated time points as given in the figures. Primary cortical neurons were treated with 1.5 µM Aβ (1-42) for indicated time points on the 6^th^ day of plating as shown in the figures.

### Plasmid constructs

shRNA specific for Sertad1 knockdown was used, based on the following sequence: 5’-CCAGACCTCCGACACCTGG-3’. A scrambled version of this sequence was used as control: shRand. For transfection in PC12 cells, shSertad1 and shRand cloned in pSiren vector backbone was used. The vector also expresses zsgreen to enable easy tracking of transfected cells. PLVTHM1 vector, which is a second-generation lentiviral vector expressing shSertad1 from the H1 promoter and GFP was used for generation of lentiviral particles. For transduction experiments, PLVTHM1 vector (EV) was used as control.

### Transfection

DNA preparation was done using the HiPure Plasmid Midiprep kit purchased from Invitrogen, Thermo Fisher Scientific (MA, USA). For survival assay, 0.5 µg of pSiren-shSertad1-zsgreen and pSiren-shRand-zsgreen plasmids were transfected on day 3 of plating primary cortical neurons and maintained for 48 hours prior to Aβ treatment. Transfection was done in 500 µl of antibiotic and serum free Neurobasal medium in 24 well plates using lipofectamine 2000. 6 hours later, medium was replaced by fresh Neurobasal medium containing B27 supplement and maintained for 72 hours prior to Aβ treatment for indicated time points (given in the figure). Similarly, 1µg of pSiren-shSertad1-zsgreen and pSiren-shRand-zsgreen plasmids were transfected on day 2 of plating PC12 cells (in the naïve condition) in OPTI-MEM medium using lipofectamine 2000. 6 hours later, cells were replenished with fresh complete medium. Priming medium supplemented with NGF was given 3-4 hours after the previous medium change. Cells were maintained for 48 hours prior to Aβ treatment.

### Animals

5xFAD (Alzheimer’s disease transgenic) mice and their non-transgenic WT littermates, C57BL6 mice were procured from The Jackson Laboratory, USA. 5xFAD mice harbour five mutations as follows: Swedish (K670N, M671L), Florida (I716V) and London (V717I) in amyloid-beta precursor protein (APP), and M146L and L286V in presenilin 1 (PS1) regulated by mouse Thy1 promoter to drive overexpression in the brain resulting in recapitulation of major pathologic features of familial Alzheimer’s disease.

### Animal housing and care

Male and female 5xFAD and WT mice were housed separately in pathogen free environments in individually ventilated cages (IVC) with sufficient bedding material. At most five mice were kept in individual cages in a controlled environment with temperatures of 22-25°C, 12-12 hour light-dark cycle, 60-70% humidity and were given access to food and water ad libitum in the animal house of CSIR-Indian Institute of Chemical Biology, Kolkata. For age-dependent experiments, male and female mice of 1 month, 6 months and 12 months each from 5xFAD and WT group were used. For all other experiments, 5 months old male and female mice from 5xFAD and WT group were employed. We have used roughly equal numbers of male and female animals to eliminate any effects of sex on phenotype. Details of number of mice used for each experiment is given in the supplementary section and in the figure legends. All experimental procedures were carried out in strict compliance with the guidelines formulated for the care and use of animals by the Institutional Animal Ethics Committee (IAEC), CSIR-Indian Institute of Chemical Biology, Kolkata.

### Lentiviral particles preparation, titre quantification and concentration

A second generation lentiviral vector PLVTHM1 expressing shSertad1 under H1 promoter at Mlu1 site was used. EV, PLVTHM1 was used as control. Lentiviral particles were produced as described previously [70]. HEK293T cells were co-transfected with PLVTHM-shSertad1-EGFP (shSertad1) or PLVTHM1-EGFP (EV), packaging plasmid, psPax2 and envelope plasmid, pMD2 using lipofectamine 2000 in 10 cm dish in OPTI-MEM medium. Lentiviral supernatant was collected post 48 hours and 72 hours of transfection. Supernatant was then centrifuged at 500g for 10 minutes to remove any dead cells and debris. To concentrate lentiviral particles were then centrifuged using Lenti-X concentrator as per the manufacturer’s protocol. Briefly, lentiviral supernatant was mixed in the appropriate ratio by gently inversion, incubated at 4°C for 30 minutes and then centrifuged at 1500 g for 45 minutes at 4°C. An off-white pellet was visible after centrifugation. Supernatant is removed carefully taking care not to disturb the pellet. Pellet was resuspended in 1/10^th^ to 1/100^th^ of the volume in Neurobasal medium, aliquoted as single use aliquots and stored at -80°C until further use. Titre was calculated as described previously [71]. Briefly, lentiviral particles are lysed using 0.2% SDS to isolate viral DNA and particle number was calculated to be 1.6×10^12^ particles/ml.

### Transduction

HEK293T cells were cultured in DMEM high glucose medium supplemented with 10% HI-FBS. 300-400 µl of lentiviral supernatant collected after 48 hours were fed in 6 well plates and samples were harvested 72 hours after transduction to assess efficacy of knockdown.

### Stereotaxic procedure

Male and female 5xFAD mice and WT mice (5 months old) were used for all experiments. Mice were anesthetized using a mixture of ketamine (60 mg/kg i.p.) and xylazine (8 mg/kg i.p.) intraperitoneally for a short time to carry out the stereotaxic injection procedures. Each mouse was placed on a stereotaxic frame (Stoelting, MO, USA). PLVTHM1-shSertad1-EGFP expressing 0.8×10^10^ lentiviral particles in 5µl were injected bilaterally in the CA1 region of the hippocampus (anterior-posterior axis: -0.2 cm, lateral axis: ±0.18 cm and dorsal-ventral axis: -0.15 cm from the bregma point). Flow rate was maintained at 0.5 µl/minute initially for 5 minutes and then at 0.2 µl/minute for 10 minutes using a worker bee syringe pump (BAS, West Lafayette, USA). After infusion at each brain hemisphere, diffusion time of 15 minutes was given to prevent any backflow of viral particles. 5xFAD and WT mice were similarly injected using concentrated lentiviral particles expressing EV.

### Behavioural experiments

Mice were divided into four groups: (1) EV-WT, (2) EV-5xFAD, (3) shSertad1-WT and (4) shSertad1-5xFAD. All behavioural investigations were carried out 14 days after bilateral stereotaxic surgery. Mice were handled and kept in the experimentation room prior to the conduction of behavioural tests to habituate them with the environment.

#### Open field test

Open field test is the standard test for measurement of anxiety like behaviour in mice. The experiment was carried out as described previously [72, 73]. The apparatus consists of an open arena of dimensions 278×236×300 mm made of plexiglass with emitters and receptors placed at equal distance along the perimeter creating an x-y grid of infrared rays. We have used the ActiTrac software (Panlab, Harvard apparatus, Barcelona, Spain) for efficient tracking of mouse trajectories. An inner square area encompassing one-third of the total arena was considered as inner zone and the remaining peripheral area was considered as the outer zone. The total distance travelled (cm), time spent in outer zone (seconds) and time spent in the inner central zone (seconds) were recorded by the software.

#### Fear conditioning test

Fear conditioning tests are routinely used to assess long-term fear based associative learning involving amygdala, hippocampus and certain cortical regions. A sound-proof chamber with steel-grid floor to deliver foot-shock is used for the experiment. The chamber is connected to a computer that has PACKWIN 2.0 software (Panlab, Barcelona, Spain) for recording the freezing behaviour in mice. Contextual and cue-dependent fear conditioning experiments were performed as described earlier [72, 73].

#### Context dependent fear conditioning

On the acquisition day, each mouse was placed in the chamber and given 2 minutes for acclimatization to the environment. Following this, mice were subjected to four trials of an unconditional stimulus (89 seconds of inter-trial interval followed by 1 second 0.4 mA foot-shock) and then a rest time of 2 minutes. Next day, mice were returned to the same chamber, but no foot-shock was given. The PACKWIN 2.0 software (Panlab, Barcelona, Spain) records percentage freezing during 8 minutes of experiment run time, which is indicative of retention of context dependent fear memory.

#### Cue dependent fear conditioning

Similar to context dependent fear conditioning, on the acquisition day, each mouse was placed in the chamber and given 2 minutes for acclimatization. The mouse was then subjected to four trials; 29 seconds of light and sound (80 dB and 2500 Hz) terminating in 1 second foot-shock, followed by 2 minutes of rest. Next day, the context was changed by attaching white sheets on the walls of the chamber. This ensures that context has no role to play in this test, and that mice only respond to light and sound cues. The protocol was run, without any foot-shock. The PACKWIN 2.0 software (Panlab, Barcelona, Spain) is used to record percentage freezing that reflects retention in cue-dependent fear memory.

#### Novel Object Recognition

This test is routinely used to measure long term recognition memory involving the hippocampus. Experiment was conducted as described previously [72, 73]. Briefly, each mouse was allowed to explore a rectangular box of dimensions 50 cm x 30 cm with two objects of same colour and shape for 5 minutes on the acquisition day. On the test day, one of the similar objects were replaced with a differently shaped object and each mouse was allowed to explore the objects for a total time of 5 minutes. Time spent near both the objects on the acquisition day and test day were recorded and tabulated. Discrimination index was calculated by the following formula: discrimination index (DI)= (TN-TF)/(TN+TF) on day 2, where TN-time spent near novel object and TF-time spent near familiar object.

#### Morris Water Maze

This test is a classic test to measure spatial and navigational learning and memory involving hippocampus. Experiment was carried out as described previously with some modifications [74]. Briefly, Morris Water Maze was setup with a hidden platform 0.5 cm below the water level (water was made opaque so that mice are unable to see the platform). The path traversed by mice in cm, time taken to reach the platform (escape latency) in seconds, number of target quadrant entries, time spent in the target quadrant in seconds was recorded by the ANY-MAZE software (Stoelting, MO, USA). Mice were allowed to swim in the maze and trained to find the hidden platform. Trials were given over three consecutive days, four trials each day. Mice were exposed to the maze in four different directions of the maze, by changing the order of directions. For example, day 1: north-east, south-east, north-west, south-east, day 2: south-west, south east, north-west, north-east and day 3: north-west, south-east, north east, south-west. Mice were removed from the maze after 90 seconds if they could not find the platform at all. On the fourth day, the platform was removed and mice were exposed to the maze to swim for 90 seconds. Time spent in the target quadrant and number of times mice enter target quadrant was recorded. Mice which spend more time in the target quadrant and keep on returning to the target quadrant have good spatial and navigational memory.

### Preparation of protein lysate

For lysis of cell and tissue sample, RIPA buffer purchased from Sigma (St. Louis, USA) along with phosphatase inhibitor (Roche, Basel Switzerland) and protease inhibitor cocktail (Takara Bio, Kusatsu, Japan) was used. Cells were scraped in lysis buffer, centrifuged at 12,000 rpm for 20 minutes at 4°C. Cortex and hippocampus from mice brain was dissected out and homogenized in prepared RIPA buffer with phosphatase and protease inhibitors to obtain a solution. The solution was then centrifuged at 12,000 rpm for 20 minutes at 4°C to remove all tissue debris. 5X sample buffer was then added to cell and tissue lysate respectively to a final concentration of 1X and then subjected to Western blotting.

### Western blot assay

30 µg protein was run on SDS-PAGE. Resolved proteins were then transferred to PVDF membrane (Merck, Darmstadt, Germany). Membrane was blocked using 5% BSA in TBS (0.15 M NaCl and 0.05 M Tris, pH 7.5). Membrane was then incubated with primary antibody incubated at 4°C overnight. Next day, membrane was washed using 0.1% Tween-20 in TBS (TBST) followed by incubation with appropriate HRP tagged secondary antibody for 1 hour at RT. Membrane was then washed with TBST prior to detection. Protein bands were visualized on iBright Imaging System (Thermo Fisher Scientific, MA, USA). Densitometric analysis of protein bands was done using NIH ImageJ software (NIH, Bethesda, USA).

### Immunohistochemistry

After completion of the experiment, at the appropriate time, mice were transcardially perfused under anaesthesia with urethane (1 mg/kg body weight in 0.9% normal saline) using ice-cold PBS and 4% w/v PFA solution. Individual mice brain was isolated and kept in 4% PFA for further fixation for 48 hours and then kept in 30% sucrose prepared in PBS for another 48-72 hours until the brain sinks stored at 4°C. 30 µm floating brain sections were taken using cryotome and stored in cryoprotectant solution (15% sucrose, in PBS) at - 20°C until further use. Sections were washed with PBS followed by permeabilization with 0.4% Triton X-100 in PBS (PBST) for 30 minutes. Sections were washed with 0.1% PBST followed by blocking with 3% goat serum in 0.1% PBST. Sections were incubated with primary antibody at the appropriate dilution overnight at 4°C. Next day, sections were washed with 0.1% PBST, followed by appropriate secondary antibody tagged with Alexa fluor prepared in blocking solution for 1-2 hours at RT. Sections were washed, nuclei were stained with Hoechst for 30 minutes at RT and washed again. Sections were then carefully mounted on glass slides using Prolong Gold Anti Fade with DAPI. Microscopic images were taken in Leica STED confocal microscope (Wetzlar, Germany). Image analysis was done using NIH, ImageJ software (NIH, Bethesda, MD, USA).

### DAB staining

DAB staining was done using VECTASTAIN ABC Kit, Peroxidase purchased from Vector Laboraties (Newark, CA, USA) as per manufacturer’s protocol. Briefly, 30 µm brain sections were collected from transcardially perfused and cryopreserved mice brain using a cryotome machine at -25°C. Free floating brain sections were washed in TBS. Sections were quenched using 3% hydrogen peroxide (H_2_O_2_) in 0.1 % TBST. Sections were then washed in TBST followed by blocking in 4% BSA prepared in 0.1 % TBST. Brain sections were then incubated with primary antibody prepared in blocking solution at the appropriate dilution overnight at 4°C. Next day, sections were washed with TBS and biotinylated secondary antibody was added and incubated at RT for 1 hour. Avidin-biotin complex is prepared in the ratio 1:1 in TBS and incubated at 37°C for 30 minutes. After secondary antibody incubation, sections were washed in TBS and then avidin-biotin complex was added and incubated for 30 minutes at RT. Sections were then washed with TBS. DAB solution was prepared in 0.1% H_2_O_2_ and then added to the brain sections and incubated till colour develops. Sections were then washed in TBS and mounted using Prolong Golf Anti Fade with DAPI for microscopic analysis. Microscopic images were taken using light microscope.

### Immunocytochemistry

Primed PC12 cells were fixed using 0.4% PFA for 15-20 minutes and then washed with PBS solution. Cells were blocked using 3% goat serum in 0.3% Triton X-100 in PBS (PBST) for 1 hour at room temperature (RT). Cells were incubated with primary antibody at appropriate dilution overnight at 4°C. Following primary antibody incubation, cells were washed with 0.3% PBST and then incubated with species matched secondary antibody tagged with Alexa fluor 546/488 for 1-2 hours at RT. Cells were washed with 0.3% PBST followed by labelling of nuclei using Hoechst for 30 minutes at RT. Cells were then washed with PBS and then mounted on glass slides using Prolong Gold Anti Fade with DAPI. Microscopic images were taken in Leica STED confocal microscope (Wetzlar, Germany). Image analysis was done using NIH ImageJ software (NIH, Bethesda, MD, USA).

### Terminal deoxynucleotidyl transferase dUTP nick end labelling (TUNEL) assay

TUNEL assay was performed using the In-situ Cell Death Detection Kit, TMR red purchased from Roche, Merck (Basel, Switzerland) as per manufacturer’s protocol. Briefly, brain sections were perfused in 4% PFA and sectioned using a cryotome. Brain sections were washed with PBS. Sections were permeabilized with 0.4% PBST for 30 minutes. TUNEL assay working solution was prepared by mixing 50 µl of Enzyme solution with 450 µl of Label solution and mixed well. Sections were labelled with the working solution at 37°C for 1 hour. Sections were washed with PBS. Sections were then labelled with Hoechst for 30 minutes at room temperature (RT) followed by washing with PBS and mounted on glass slides using Prolong Gold Antifade with DAPI for microscopic analysis. Microscopic images were taken in Leica STED confocal microscope (Wetzlar, Germany). Percentage of apoptotic cells was determined by counting the number of TUNEL+ cells per total cell number expressed as percentage.

### Golgi-cox impregnation and staining

Golgi-cox staining was performed as described previously [75]. Mice brains were isolated after 21 days of stereotaxic surgery were immersed in an impregnation solution, mixture of five parts by volume solution A (5% solution of potassium dichromate), five parts by volume solution B (5% mercuric chloride solution), four parts by volume solution C (5% potassium chromate solution) and ten parts distilled water and stored in the dark for 24 hours at RT. Next day, solution is replaced with fresh impregnation solution and kept in the dark at RT for 2-3 weeks. Mice brains were then transferred to solution D (30% sucrose in PBS) for cryopreservation at 4°C. Once the brain sinks to the bottom of the tube, 75 µm brain sections were collected using cryotome at -25°C on clean glass slides. Sections were washed with distilled water, then immersed in 20% ammonia solution for 10-15 minutes in the dark for staining. Brain sections were then dehydrated using 50%, 75% and 100% ethanol for 5 minutes each. Lipids were removed by two washed in xylene for 10 minutes each. Sections were mounted using DPX reagent and allowed to dry for 4-5 days before imaging. Microscopic images were taken using Leica STED confocal microscope (Wetzlar, Germany). Spine density analysis was done as described previously [76]. Any small protrusion was considered a dendritic spine. 50-80 dendritic stretches were analysed per experimental group in the cortex and CA1 region of hippocampus respectively using Adobe Photoshop (San Jose, California, USA).

### Statistical analysis

All data analysis and graph preparation were done using the Graphpad Prism software (v9.0.0 (121), Graphpad Inc, USA). Statistically significant difference between two groups was calculated using two-tailed unpaired t-test, while significant differences between more than two groups was done using one-way ANOVA followed by Tukey’s multiple comparison post-hoc test except mentioned otherwise. All data are represented as mean ± standard error of mean (SEM) and p<0.05 was considered as statistically significant.

## Abbreviations

Aβ: amyloid-β
AD: Alzheimer’s disease
AL: autolysosome
ALN: autophagy-lysosome network
AV: autophagic vacuoles
ATG: autophagy related
CDK4: cyclin dependent kinase 4
CTCF: corrected total cell fluorescence
E2F1: E2F Transcription Factor 1
EGFP: enhanced Green Fluorescent Protein
EV: empty vector
FAD: familial Alzheimer’s disease
ICC: immunocytochemistry
IHC: immunohistochemistry
LAMP1: lysosomal associated membrane protein 1
LC3: Microtubule-associated protein 1A/1B-light chain 3
LTP: long term potentiation
MAP2: Microtubule Associated Protein 2
NGF: Nerve Growth Factor
NOR: Novel Object Recognition
PAS: phagophore assembly site
PP2A: protein phosphatase 2A
Rb: retinoblastoma
SEM: standard error of mean
Sertad1: Serta-domain containing 1
SMAD1: SMAD Family Member 1
SQSTM1: sequestosome 1
TUNEL: Terminal deoxynucleotidyl transferase dUTP nick end labelling
XIAP: X-linked inhibitor of apoptosis protein.

## Acknowledgements

We would like to thank Mr. Sounak Bhattacharya from the Central Instrument Facility (CIF), CSIR-IICB, Kolkata for his contributions to carry out Leica Sp8 STED confocal microscopy image acquisition. We thank Dr. Priyankar Sanphui for valuable discussions. We thank Dr. Debabrata Biswas for providing psPAX2 and pMD2 plasmids for lentivirus preparation.

## Author contributions

N.A. and S.C.B conceived and designed the study. S.C.B. supervised the study, acquired funding for the experiments and provided all the necessary resources for carrying out the work. N.A. did all the biochemical, immunofluorescence and behavioural studies. N.A. and K.G. performed animal stereotaxy for knockdown of Sertad1 and immunohistological experiments and Golgi staining. N.A. analysed all the data and wrote the original draft of the manuscript. S.C.B. reviewed and edited the manuscript and contributed to writing of the paper.

## Ethics statement

All animal studies were carried out in accordance with the guidelines formulated bythe Committee for the Purpose of Control and Supervision of Experiments on Animals (Animal Welfare Divisions, Ministry of Environments and Forests, Govt. of India) with approval from the Institutional Animal Ethics Committee (IAEC). We have consulted the ARRIVE guidelines for the relevant aspects of animal studies.

