## Supplementary Materials for "Sertad1 is elevated and plays a necessary role in synaptic loss, neuron death and cognitive impairment in a model of Alzheimer’s disease"

**Supplementary information**

To check if Sertad1 has any role in autophagy modulation in cellular model of AD, we utilized primed PC12 cells treated with oligomeric Aβ. We transfected shSertad1 or shRand in cultured PC12 cells and maintained in NGF containing medium for differentiation of cells. After maintaining cells for 72 hours, they were treated with oligomeric Aβ for 16 hours. Post-treatment, cells were fixed with 4% PFA in 1x PBS. We then assessed LC3 and p62 levels and quantified the number of puncta per cell respectively. We found that shSertad1 transfected cells showed lower LC3 staining (Fig. S1A and B) and fewer number of LC3 positive puncta per cell (Fig. S1A and C). Further, shSertad1 transfected cells showed lower expression of p62, (Fig. S1E and F) and fewer number of p62 puncta per cell (Fig. S1E and G). Our results show that Sertad1 knockdown cells show lower levels of autophagy markers in agreement with the data obtained in in-vivo experiments.

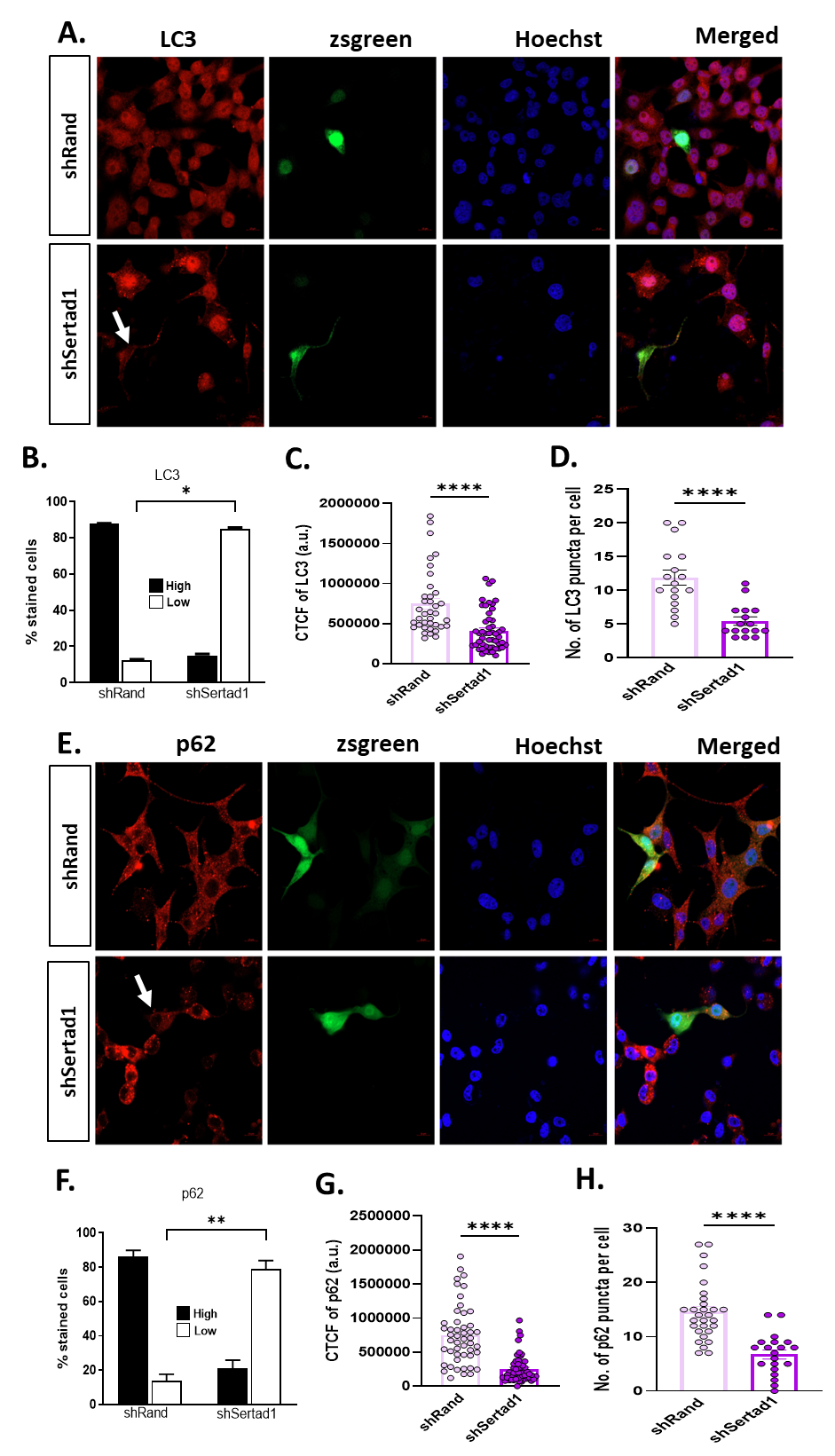

**Figure S1:** PC12 cells were transfected with pSIREN-shSertad1-zsgreen (shSertad1) and pSIREN-shRand-zsgreen (shRand), primed using NGF and maintained for 72 hours followed by treatment with 3µM Aβ for 16 hours. (A) Representative images of transfected PC12 cells upon immunostaining with LC3 antibody. Left to right: first panel shows expression of LC3 (red), second panel shows zsgreen (green) expression, third panel depicts nuclei stained with Hoechst (blue) and fourth channel shows merged image, taken at 63X magnification. (B) Percentage-stained cells indicate the number of transfected cells (green) that show high (high or equal intensity as compared to the non-transfected neighbouring cell) or low (low intensity as compared to the neighbouring non-transfected cell) after Aβ treatment. Graphical representation of CTCF of LC3 (C) and number of LC3 puncta per cell (D). (E) Representative images of transfected PC12 cells upon immunostaining with p62 antibody. Left to right: first panel shows expression of p62, second panel shows zsgreen expression, third panel depicts nuclei stained with Hoechst and fourth channel shows merged image, taken at 63X magnification. (F) Percentage-stained cells indicate the number of transfected cells (green) that show high (high or equal intensity as compared to the non-transfected neighbouring cell) or low (low intensity as compared to the neighbouring non-transfected cell) after Aβ treatment. Graphical representation of % stained cells is depicted. 50 cells were counted per experiment. Graphical representation of CTCF of p62 (G) and number of p62 puncta per cell has been depicted (H). Approximately 15-20 cells were analyzed per group per experiment and 3 independent experiments were performed. Data represent Mean±SEM of three independent experiments. Statistical analysis was done using One-way ANOVA, Tukey’s post-hoc analysis. Asterisks denote statistically significant differences between indicated groups; *p<0.05, **p<0.01 and ****p<0.0001.

**Supplementary Table 1:**

| **Experiment** | **GROUPS** | **BATCH 1**  **(FEMALES)** | **BATCH 2**  **(MALES)** | **TOTAL** |
| --- | --- | --- | --- | --- |
| Locomotion | EV Control | 5 | 5 | 10 |
|  | EV TG | 3 | 4 | 7 |
|  | shSertad1 Control | 4 | 4 | 8 |
|  | shSertad1 TG | 4 | 3 | 7 |
| NOR | EV Control | 5 | 5 | 10 |
|  | EV TG | 3 | 4 | 7 |
|  | shSertad1 Control | 4 | 4 | 8 |
|  | shSertad1 TG | 4 | 3 | 7 |
| Cue and context dependent fear conditioning | EV Control | 5 | 5 | 10 |
|  | EV TG | 3 | 4 | 7 |
|  | shSertad1 Control | 4 | 4 | 8 |
|  | shSertad1 TG | 4 | 3 | 7 |
| EPM | EV Control | 5 | 5 | 10 |
|  | EV TG | 3 | 4 | 7 |
|  | shSertad1 Control | 4 | 4 | 8 |
|  | shSertad1 TG | 4 | 3 | 7 |
| Morris Water Maze | EV Control | 7 | 7 | 14 |
|  | EV TG | 4 | 4 | 8 |
|  | shSertad1 Control | 3 | 4 | 7 |
|  | shSertad1 TG | 4 | 3 | 7 |
